# Ribonucleotide reductases recapitulate biogeographic patterns within virioplankton according to ocean biogeochemistry

**DOI:** 10.64898/2026.02.11.705308

**Authors:** Amelia O. Harrison, Ryan M. Moore, Barbra D. Ferrell, Shawn W. Polson, K. Eric Wommack

## Abstract

Many of the important biogeochemical implications of marine viruses focus on the activity of lytic viruses. It is generally thought that lytic viral activity contributes to maintaining bioavailable nutrient pools and controlling host community composition through specific infection and lysis processes. Among lytic viruses, genome replication is under stringent selection pressure as advantages in replication speed and fidelity can influence viral fitness. Deoxyribonucleotides (dNTPs) are the precursor substrate for DNA synthesis and ribonucleotide reductase (RNR) is the only known enzyme capable of reducing ribonucleotides into dNTPs. Thus, for a virus, encoding an RNR gene provides control over dNTP production. Overall, distributional patterns of the RNR-encoding virioplankton community mirrored that seen for total virioplankton throughout the global ocean. A majority of RNR-encoding virioplankton (∼66%) carried the Class II enzyme and most of these carried the monomeric (NrdJm) gene. This was significant as NrdJm utilizes triphosphate ribonucleotides as opposed to the diphosphate ribonucleotides used by Class I and Class II dimeric RNRs. The distribution and frequency of Class I RNRs followed the concentration of required iron and manganese co-factors. Oceanic virioplankton encompassed a high diversity of Class I and II RNRs with the discovery of new phylogenetic clades not previously observed among viruses or within cellular life. Representing ∼10% of all virioplankton populations, RNR-encoding viral populations demonstrated surprising fidelity with the major biogeographical features defining oceanic ecosystems.

## Introduction

Viruses are critical players in marine ecosystems. In particular, they are known to influence biogeochemical cycling via infection of microbial hosts (Fuhrman 1999), with the viral shunt being perhaps the most famous example (Shiah et al. 2022; Locke et al. 2022; Wommack and Colwell 2000). Additionally, viruses can affect the flow of nutrients through the environment via the termination of algal blooms (Laber et al. 2018) and facilitation of horizontal gene transfer (Lindell et al. 2004).

Virioplankton can also take more active roles in shaping biogeochemical cycles. Preferential packaging of certain elements in viral particles changes the relative abundance of elements in the environment (Jover et al. 2014), and cyanophages have been shown to carry out photosynthesis and inhibit carbon dioxide fixation when infecting their cyanobacterial hosts (Puxty et al. 2016). The latter example is made possible by cyanophage encoding metabolic genes (those involved in photosynthesis and photosynthetic electron transport). While these examples of phages altering basal cellular metabolism are compelling, during active infection all phages, indeed all viruses, share a common need for available deoxyribonucleotides supporting viral genome replication, which are generally kept at low levels in bacterial cells through tight control of ribonucleotide reductase outside of the S phase or DNA damage events (Torrents 2014; Mathews 2006; Nordlund and Reichard 2006).

Ribonucleotide reductase (RNR) is a nucleotide metabolism protein commonly encoded by lytic dsDNA phages and is solely responsible for producing deoxyribonucleotides (dNTPs) *de novo*, the rate-limiting step of DNA synthesis. For most viruses, simply recycling dNTPs from the host genome would not provide sufficient material for supporting observed burst sizes (Kirzner et al. 2016; Mahmoudabadi et al. 2017). Thus, the need for producing deoxyribonucleotides sufficient for phage replication places evolutionary pressure on a phage to carry an RNR. In fact, a previous survey of phage protein content found RNR to be present in 17.4% of all dsDNA phages (Wommack et al. 2015).

Ribonucleotide reduction requires radical chemistry. RNRs are divided into three main classes, some with multiple subclasses, based on the biochemical method used for radical production (Figure 1). Thorough descriptions of the different RNR types are available elsewhere (Ruskoski and Boal 2021; Harrison et al. 2019; Torrents 2014; Kolberg et al. 2004). Briefly, Class I RNRs are oxygen-dependent and most use a di-metallic cofactor consisting of iron, manganese, or both to produce a radical (one subclass has evolved to be metal-free and instead uses a dihydroxyphenylalanine (DOPA) radical). Class I subclasses are divided based on the identity of the cofactor. Class II RNRs are oxygen-independent and produce a radical by cleaving adenosylcobalamin (coenzyme B_12_). Class II subclasses are divided based on quaternary structure (dimeric (Class IId) or monomeric (Class IIm)) and the phosphorylation status of the ribonucleotide substrate. Class III RNRs are oxygen-sensitive and require an iron–sulfur cluster for radical production. Only two RNRs, Class IIm and Class III utilize a triphosphate ribonucleotide substrate (dNTPs), all others use a diphosphate substrate (dNDPs). All RNR classes and subclasses share a common ancestor (Lundin et al. 2015) and are, so far, monophyletic (Burnim et al. 2022). A fourth group of RNRs has previously been referred to as Cyano SP (Harrison et al. 2019) and Cyano II (Sakowski et al. 2014) because its identified members consist of cyanosipho- and podoviruses. This group was recently proposed as a novel class of RNRs, Class ø (Burnim et al. 2022), but is yet to be biochemically characterized. Here, we mainly refer to this group as Cyano SP.

**Figure 1.**
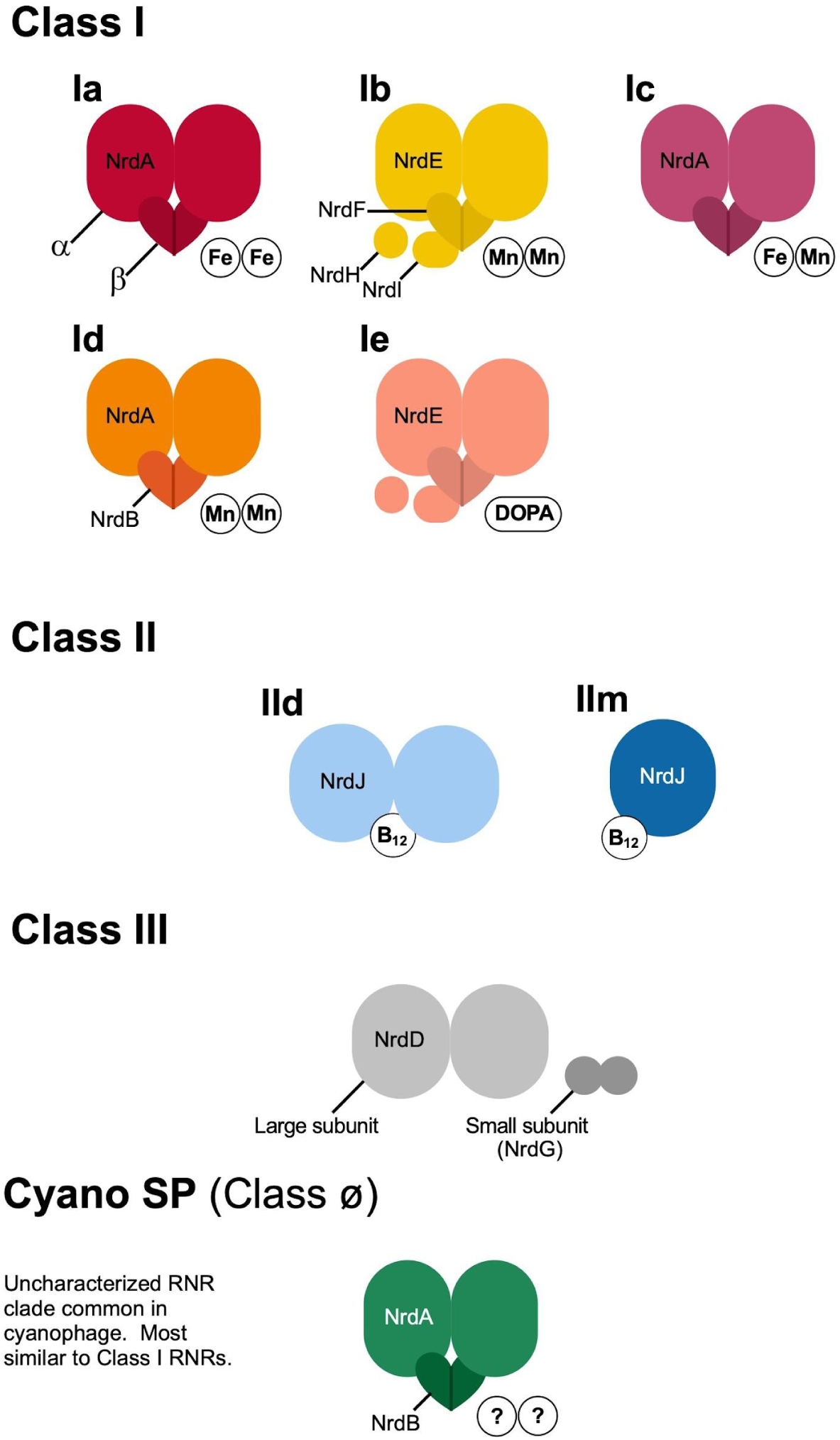
Schematic of ribonucleotide reductase classes. Subclasses are designated according to their quaternary structure of multimers along with metal (Fe, Mn), vitamin (B12), or amino acid (dihydroxyphenylalanine (DOPA)) co-factors.

Class I and II RNRs are commonly found in the genomes of marine viruses (Wu et al. 2023; Wommack et al. 2015; Sakowski et al. 2014; Dwivedi et al. 2013), while Class III RNRs are virtually absent from the pelagic ocean because of the presence of oxygen (Dwivedi et al. 2013). Class I and II RNRs require trace metals either as cofactors or as metal centers within their cofactors (Grinberg et al. 2019; Blaesi et al. 2018; Nordlund and Reichard 2006; Morel and Price 2003). In the oceans, trace metals are often limiting nutrients (Lohan and Tagliabue 2018; Browning et al. 2017; Sunda 2012). Distribution of most trace metals in the ocean changes with depth and geographical location reflecting changes in input sources and chemical reactions that influence valence state (Lohan and Tagliabue 2018; Sunda 2012). Because RNRs may influence viral fitness and are reliant upon differentially distributed trace nutrients, we hypothesized that marine viruses carrying different types of RNR would show different biogeographical patterns and that these patterns would mirror those of the trace metals required by RNRs.

The potential connection between RNR type and ecological distributions of viral populations make RNR an intriguing marker gene candidate. Currently, most marker genes target specific viral populations (Finke and Suttle 2019; Needham et al. 2017; Adriaenssens and Cowan 2014). However, RNR occurs within the genomes of diverse viruses infecting a broad cross section of microbial hosts (Iranzo et al. 2016; Sakowski et al. 2014). Thus, RNRs have the potential to be informative on the population and community levels. RNRs may also be indicative of phage biology, as they are strongly tied with phages having a virulent lifestyle (Nasko et al. 2018; Wommack et al. 2015; Sakowski et al. 2014). Finally, given its central role in DNA genome replication, RNRs are of interest within evolutionary biology (Lundin et al. 2010; Wächtershäuser 2006) and cancer research (Greene et al. 2020; Nordlund and Reichard 2006), and are thus abundant in sequence databases and scientific literature. Therefore, we believe that RNR is a valuable marker gene for understanding the ecological behavior of environmental viral populations.

RNR sequences were obtained from the Global Ocean Virome (GOV) 2.0 dataset to test whether RNR subclass drives virus ecological distributions and demonstrate the utility of RNR as a marker gene for exploring the virioplankton community ecology. The GOV 2.0 dataset contains 145 viromes from globally distributed marine sites and is accompanied by extensive metadata, making it useful for ecological study. Additionally, the total viral community from GOV 2.0 has previously been analyzed (Gregory et al. 2019); enabling comparison of RNR-encoding viral populations to prior virioplankton community analyses.

## Materials and Methods

### Data Sources

The GOV 2.0 dataset (Gregory et al. 2019), which contains 145 viromes from *Tara* Oceans, *Tara* Oceans Polar Circle (TOPC), and the Malaspina expedition, was used for exploring the diversity and ecology of RNR-encoding viruses in the world’s oceans. These viromes are globally distributed and were sampled largely from four depth zones: surface (5 m depth; n = 60), deep chlorophyll maximum (DCM; n = 39), mesopelagic (n = 26), and bathypelagic (n = 13). Samples were taken at the mixed layer depth on the rare occasion that a DCM could not be identified (n = 6).

GOV 2.0 viral populations ≥5 kb nucleotides or circular were downloaded from the CyVerse Data Commons (https://datacommons.cyverse.org/browse/iplant/home/shared/iVirus/GOV2.0). Environmental data associated with each of the viromes was downloaded from the GOV 2.0 paper supplement (Gregory et al. 2019). Viral population abundances and Malaspina JGI paths were downloaded from the MAVERIC-Lab bitbucket (https://bitbucket.org/MAVERICLab/gov2/downloads/). Relevant details regarding the GOV 2.0 data collection and processing can be found in the <u>Supplementary Methods</u>.

### Identifying RNRs within GOV 2.0 viral populations

ORFs were predicted and translated for GOV 2.0 viral populations ≥5 kb or circular using Prodigal v2.6.3 with option -meta (Hyatt et al. 2010). A homology search using the resulting peptides was performed against Class I α, Class I β, Class II, and Class III large subunit peptide sequences from the RNRdb (Lundin et al. 2009). RNRdb protein sequences were clustered using CD-HIT v4.6 to remove exact and sub-string matches (Fu et al. 2012; Li and Godzik 2006). The homology search was performed using MMseqs2 v5ae55 easy-search with the highest sensitivity setting (-s 7) (Steinegger and Söding 2017).

Verification steps were performed on the resulting putative RNR peptide sequences using PASV v2.0.2 (Moore et al. 2021). PASV eliminated bycatch from the sensitive homology search and ensured that only functional RNRs were used for data analysis (there is at least one recorded case of a viral RNR with mutated active sites that now performs a function other than ribonucleotide reduction (Lembo et al. 2004)). The pasv-msa command was used for aligning each query sequence individually with a set of reference sequences, and partitioning them based on amino acid residues in specified positions and whether they spanned a specified region of interest. Clustal Omega v1.2.4 was used by PASV as the aligner for all RNR types (Sievers et al. 2011).

Two sets of PASV reference sequences for RNR verification (one for Class III RNR large subunit sequences, and one for both Class I α subunit and Class II sequences) used sequences from the RNRdb in accordance with PASV’s recommended best practices. A single reference set was suitable for Class I α and Class II RNR sequences because they share many conserved residues (Moore et al. 2021; Lundin et al. 2015; Sakowski et al. 2014). Residue positions and identities and regions of interest used for verification of each RNR type are available in Supplementary Tables S1 and S2. PASV profiles and reference sets underwent testing using RNRdb sequences prior to their use here (data not included).

Putative RNR sequences resulting from the homology search were run through both PASV profiles, resulting in two sets of sequences: putative Class III large subunit sequences and putative Class I α subunit/Class II sequences.

### RNR curation and classification

GOV 2.0 RNR sequences verified by PASV were imported into Geneious v10.2.6 (https://www.geneious.com) for manual curation and classification. First, sequences were aligned with appropriate PASV reference sequences (e.g., putative Class III sequences were aligned with a subset of RNRdb Class III references) using the MAFFT Geneious plug-in v7.450 on the AUTO setting with scoring matrix BLOSUM62 (Katoh and Standley 2013). Any sequences not containing the conserved residues or spanning the ROI used for PASV validation were removed from further analysis. All 202 sequences passing the Class III profile were removed in this step, with 199 of them determined to be Class II sequences that passed both PASV profiles.

Inteins were manually removed from the remaining Class I α and Class II sequences in Geneious using alignment gaps and conserved residues to determine intein start and stop positions (Wang et al. 2022; Shah and Muir 2014). All sequences were then trimmed to the region of interest to remove evolutionarily mobile domains (e.g., ATP cones (Aravind et al. 2000)), and non-homologous domains (e.g., the Class II B_12_ binding domain) that could interfere with phylogenetic classification.

Trimmed sequences underwent phylogenetic classification as described in (Harrison et al. 2019). For all classification steps, sequences were aligned with appropriate references from the RNRdb using the MAFFT plug-in as described above and phylogenies were inferred using the FastTree Geneious plug-in v2.1.11 with default settings (Price et al. 2010). RNRdb reference sequences were previously curated, trimmed, and clustered as described in (Harrison et al. 2019). A phylogenetic tree was used to separate Class I α subunit and Class II sequences prior to further classification. A Class I α subunit tree was used to divide sequences into RNRdb clades (Supplementary Figure S1), with sequences from the Cyano M and Cyano SP (Class ø) clades (Harrison et al. 2019) also being identified. A Class II phylogenetic tree was used to divide sequences into monomeric and dimeric subclasses (Supplementary Figure S2).

### Validating a Novel RNR clade

During classification of GOV 2.0 RNRs, there was a large clade (269 sequences) of Class I α RNRs that did not fall within an existing RNRdb or cyanophage clade. To determine the subclass identity of the clade, contigs containing the novel Class I α RNR sequences were manually examined for the presence of Class I β subunits, as the primary sequence structure of Class I β subunits can hold clues as to Class I RNR subclass identity (Harrison et al. 2019). As Class I α and β subunits generally occur next to each other in viral genomes (Harrison et al. 2019; Dwivedi et al. 2013), nearby proteins to Class I α sequences were checked first. Putative Class I β sequences were identified by manually aligning those peptide sequences neighboring novel Class I α sequences with known Class I β sequences in Geneious. Identified putative Class I β sequences were aligned against reference sequences using the MAFFT Geneious plug-in with default settings (Harrison et al. 2019), and sequences were checked for the presence of the immutable active sites W48 and Y357 (*E. coli* numbering), as well as the presence of metal-binding residues as in (Harrison et al. 2019). Sequences that met these requirements were then checked for the identity of the residue in position 123 (*E. coli* numbering) to determine subclass membership of the Class I α and β sequences.

#### Ecological analysis

Viral populations with validated Class I α or Class II RNRs were used for ecological analysis. Populations with more than one RNR on the representative contig (13 contigs out of 18,776) were removed prior to this analysis. All ecological analyses were performed using R v4.1.1 (R Core Team 2024) and RStudio v2023.0.3.0 (Posit team 2024).

### RNR-encoding viral population abundances

#### Filtering

Average read depths of viral population contigs ≥5 kb or circular with validated RNR sequences were used as a proxy of RNR-encoding virus abundance. Viral populations with no reads mapping were removed from analysis. Samples 155_SUR and 189_SUR were excluded from further analysis; reads from sample 155_SUR mapped to only one RNR-encoding viral population, and no reads from sample 189_SUR mapped to any of the RNR-encoding viral populations. Gregory et al. also excluded sample 155_SUR from much of their analysis.

Because physicochemical data was comparatively limited in the Malaspina study, these samples were excluded in some analyses.

#### Compositional data analysis preparation

Zeros in the RNR-encoding viral population abundance tables were replaced using the square root Bayesian multiplicative method with the ‘cmultRepl’ function from R package zCompositions v1.3.4 (Palarea-Albaladejo and Martín-Fernández 2015). Following zero replacement, abundances were centered log-ratio (CLR) transformed using the function ‘clr’ from R package compositions v2.0-1 (van den Boogaart and Tolosana-Delgado 2008).

### Environmental data preparation

#### Data imputation

Analysis of RNR-encoding viral populations in relation to measured physicochemical parameters was limited to the *Tara* Oceans and TOPC expeditions because of the comparatively limited scope of similar data from the Malaspina expedition. However, many *Tara* Oceans and TOPC expeditions samples were missing physicochemical data. Missing values were imputed for nitrate, nitrite, nitrate/nitrite, phosphate, silicate, chlorophyll a, POC, PIC, temperature, oxygen, and salinity using Multiple Imputation by Chained Equations from the R package mice v3.13.0 (van Buuren and Groothuis-Oudshoorn 2011). The variables PAR1, PAR8, and PAR30 also contained missing data, but at a much greater amount than other variables, so data was not imputed for these variables and these were excluded from downstream analyses.

#### Data transformation

Physicochemical parameters with right-skewed distributions were transformed using a square root transformation to dampen the effects of outliers and to normalize their distributions ahead of statistical analysis. Transformed variables included ammonium, nitrate, nitrite, phosphate, silicate, POC, PIC, and chlorophyll a.

### Spatial data

In the oceans, the distribution of viruses and other plankton are highly dependent on the currents (Richter et al. 2022; Brum et al. 2015). As a result, GOV 2.0 samples connected by ocean currents were likely autocorrelated, which could severely affect statistical analyses. Spatial autocorrelation was tested for in the RNR-encoding viral populations and physicochemical data from the *Tara* Oceans and TOPC expeditions. Spatial models were built to account for autocorrelation in statistical testing. Bathypelagic samples from the Malaspina expedition were excluded from analyses that incorporated spatial data because they could not be accounted for in our model. Details on the initial testing of parameters for autocorrelation and the subsequent creation of spatial models can be found in the supplementary methods.

### Community structure analysis

#### Exploratory analysis

Principal component analysis (PCA) was performed directly on the CLR-transformed viral population abundance table using the R package biplotr v0.0.13 (https://github.com/mooreryan/biplotr).

Euclidean distances between samples were calculated based on the CLR-transformed viral population abundance data. Hierarchical clustering was then computed using Ward’s method with the Aitchison distance. The resulting dendrogram was colored and customized using Iroki (Moore et al. 2020).

#### Community structure and the environment

Variation partitioning was performed using the function ‘varpart’ in the R package vegan v2.5-7 (Oksanen et al. 2007). Variation was partitioned among spatial structure, continuous physicochemical variables, and ecological regions. Spatial structure was represented by significant eigenvectors from the spatial weights matrix. Measured physicochemical variables, which included imputed values, were centered and scaled prior to analysis using the function ‘scal’ within the base R software package. Ecological regions included Longhurst province, GOV 2.0 ecological zone, and depth of sampling. Ecological region variables were transformed into dummy variables prior to testing using R package fastDummies v1.6.3 (Kaplan 2020). A proportional Venn diagram was created to display variation partitioning results using the R package eulerr v7.0.0 (Larsson and Gustafsson 2018).

Following variation partitioning, each partition was tested for significance by conducting redundancy analysis (RDA) for each partition and testing the RDA for significance. RDAs were performed using the function ‘rd’ from vegan. Significance tests were performed using function ‘anova.cc’ from vegan with default settings, which performs an anova-like permutation test with 999 permutations.

Significant differences between the RNR-encoding virus communities among sample environmental groups and physiochemical parameters were determined using perMANOVA tests. These tests were conducted by performing partial distance-based redundancy analysis tests using the function ‘dbrd’ from vegan which tests the significance of individual variables on community composition, followed by significance tests using function ‘anova.cc’ from vegan with 199 permutations. R-squared values were adjusted for multiple testing using ‘RsquareAdjust’ from vegan.

### RNR types and the environment

#### Generalized additive models

Generalized additive model smooth plots were used for examining the distributions of different RNR subclasses and assessing their relationships with iron measurements. All plots were created using ‘geom_smooth’ from ggplot2 v3.4.1 using cubic splines (bs = “cs”) for smoothing with 95% confidence intervals. RNR subclass Ib-containing viral populations were excluded from this analysis because there were too few for reliable analysis (n = 15). Other RNR subclasses were plotted against iron, latitude, and depth. Iron concentrations were predicted for epipelagic *Tara* Oceans samples using two biogeochemical models developed by a leading group of ocean trace metals researchers in conjunction with the *Tara* Oceans team (Caputi et al. 2019). Predictions shown here were generated by the PISCES model. The predicted iron concentrations are considered to be more reliable than the iron measurements taken during the *Tara* Oceans expedition (Caputi et al. 2019).

### Maps

Maps were created in R using ggplot2 v3.4.1. Other packages used were rgdal v1.6-7 (Bivand et al. 2023), mapproj v1.2.11 (Brownrigg et al. 2023), maps v3.4.1 (Becker et al. 2022), sf v1.0-9 (Pebesma 2018), sp v1.6-0 (Pebesma and Bivand 2005), tmap v3.3-3 (Tennekes 2018), and raster v3.6-14 (Hijmans 2023). Shapefiles were used to add ocean currents (Hamilton 2018), landmasses (Natural Earth), and latitude and longitude lines (Natural Earth). The Lambert azimuthal equal-area projection was used for maps centering on the Arctic. The Robinson projection was used for all other maps. Arrows to indicate the directionality of cross-equatorial flow at the time of sample collection were also added manually.

Some manual modification of ocean currents was performed, including the removal of currents not near sample locations (e.g., the Gulf Stream) and the addition of local currents that were not present in the shapefile (e.g., the Red Sea Eastern Boundary Current). In one rare case, a modification was made to reverse a current (the Somali Current) as compared to the shapefile based on the date of sample collection. All current additions and modifications to existing currents were based on scientific literature and are summarized in Supplementary Table S3.

## Results

### RNR Identification and Curation

RNRs were identified within GOV 2.0 viral populations (contigs ≥ 5 kb or circular sequences) using MMseqs2 for homology searching. Subsequently, putative RNRs were verified as having critical active site residues using the protein active site validation (PASV) tool (Moore et al. 2021). PASV identified 18,726 viral protein sequences as Class I α or Class II RNR. Of these, 18,717 spanned the PASV region of interest (ROI) and had all required conserved residues upon visual inspection in a multiple sequence alignment. Eight sequences contained inteins. Thirteen viral population contigs contained two RNRs and were removed from analysis as potential chimeric assemblies, leaving 18,691 sequences, representing 9.6% of the GOV 2.0 ≥5 kb viral populations, for analysis. Of these, 18,578 viral populations were used for ecological analysis (113 were filtered out because no reads mapped).

### Phylogenetic diversity of marine viral RNRs

Class II RNRs made up nearly two-thirds (∼66%) of the RNR sequences recovered, with the monomeric subclass (Class IIm) alone making up almost half (∼49%) of all sequences. Just under a third (∼32%) of the RNR sequences belonged to Class I, with subclass Id and the uncharacterized RNRdb (Lundin et al. 2009) clade NrdAk together making up almost three quarters of the Class I sequences. Cyano SP RNRs (Class ø) made up almost two percent of the total RNR sequences. No Class III RNRs were identified.

While GOV 2.0 viral populations contained most of the described RNR Class I and Class II subclasses (Supplementary Figures S1 and S2), no sequences belonging to Class I subclasses Ic or Ie were identified and only 15 subclass Ib sequences were found (Table 1). All other Class I subclasses were well-represented, though some Class I RNRdb clades were absent (NrdAh, NrdAn, NrdAq) or found in very low numbers (NrdAm, NrdAz). RNRdb clade NrdAk was analyzed individually and not grouped into any subclass because the clade has no biochemically characterized representatives (Grinberg et al. 2019), groups away from subclass Ia clades on a phylogenetic tree (Supplementary Figure S1), and had enough viral population members (n = 1942) to be analyzed separately.

**Table 1.** Class I α subunit and Class II RNRs retrieved from GOV 2.0 by type.

| Class | Subclass | RNRdb Clade* | Sequences |  |
| --- | --- | --- | --- | --- |
|  |  |  | No. | % of total |
| I |  | Ae | 162 | 0.87 |
|  |  | Ag - non-Cyano M** | 923 | 4.93 |
|  | Ia | Ag - Cyano M** | 303 | 1.62 |
|  |  | Am | 5 | 0.03 |
|  |  | Aza | 3 | 0.02 |
|  | Ib | Eb | 15 | 0.08 |
|  | Unknown, Ic-like | _*** | 269 | 1.44 |
|  | Id | Ai | 2438 | 13.03 |
|  | Unknown | Ak | 1942 | 10.38 |
| II | II dimeric | Multiple | 3149 | 16.82 |
|  | II monomeric | Jm | 9140 | 48.83 |
| Cyano SP (∅) | - | _*** | 368 | 1.97 |
| *Total* |  |  | 18717 | 100.00 |

\* RNRdb (Lundin et al. 2009) phylogenetic clades and subclasses detailed in Harrison et al. 2019.
\*\*Cyano M RNRs are members of the NrdAg RNRdb clade, but were analyzed separately.
\*\*\*Not currently defined or present in the RNRdb.

#### A novel RNR clade

An unknown GOV 2.0 RNR clade, which formed a clade lacking any RNRdb reference sequences, was closest to the NrdAz RNRdb clade, which contains sequences belonging to both subclass Ia and subclass Ic (Supplementary Figure S1). These subclasses cannot be reliably distinguished based on the α subunit sequence alone, so the corresponding β subunit sequences were examined. Putative Class I β subunits on contigs with α subunits belonging to the unknown clade did show one classic sign of a subclass Ic β subunit in that they lacked the tyrosine residue (Y123 in *E. coli*) that harbors the stable protein radical and is conserved in subclasses Ia, Ib, Id, and Ie (Blaesi et al. 2018; Cotruvo et al. 2013; Nordlund and Eklund 1993). However, the alternate residue in that position (isoleucine or valine) was not consistent with subclass Ic reference sequences (phenylalanine or leucine). The group was therefore kept separate for ecological analysis and is referred to as subclass Ic-like.

### Community composition of RNR-encoding viruses

RNR-encoding viral communities in global marine samples were compared based on sample abundances of GOV 2.0 viral populations (95% clusters of viral contigs). Principal components analysis (PCA) performed on the CLR-transformed community abundance matrix showed that samples separated largely based on the ecological regions defined by Gregory et al. (Figure 2). Arctic and non-Arctic samples separated along PC1, which explained 16.4% of the variation.

**Figure 2.**
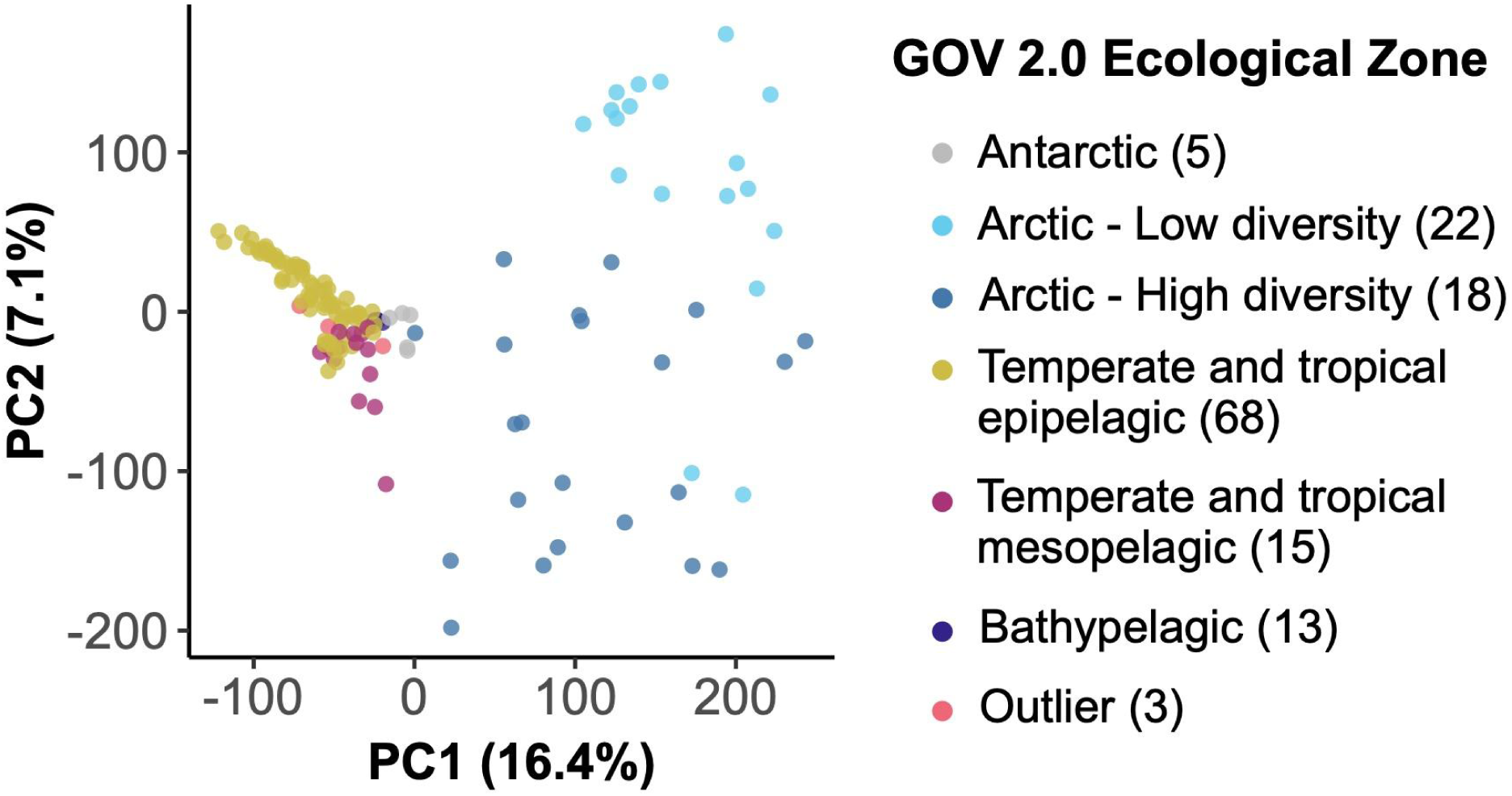
Principal components analysis (PCA) of GOV 2.0 samples based on the center log ratio-transformed abundances of RNR-encoding viral populations in each sample. Points (samples) are colored by the ecological zone of the sampling station as defined in the original GOV 2.0 paper (including samples labeled as “Outliers”). The number of samples in each ecological zone is indicated in the legend.

Temperate and Tropical Epipelagic, Temperate and Tropical Mesopelagic, Bathypelagic, and Antarctic samples also showed some separation along PC1. PC2 explained 7.1% of the variation in community structure and was driven by the separation between Arctic Low and High diversity samples. The three samples that were categorized as “Outliers” by Gregory et al. grouped with similar sample types.

A dendrogram based on the same community abundance matrix showed relationships between samples more clearly (Figure 3). The dendrogram showed 10 main clusters: Arctic High Diversity, Arctic Low Diversity, Red & Arabian Seas, Indian Ocean, Temperate and Tropical Mesopelagic, South Pacific, Mediterranean, South Atlantic, Antarctic & Upwelling, and Bathypelagic. The Arctic clusters contained both epi- and mesopelagic samples from the region. In contrast, samples from non-polar regions largely formed separate epipelagic and mesopelagic clusters. The Antarctic & Upwelling cluster contained all of the Antarctic samples (except Station 85 MES) and samples from major upwelling zones, such as the Peruvian and Benguelan coasts (Ohde and Dadou 2018; Tarazona and Arntz 2001; Nelson and Hutchings 1983; Brink et al. 1983). Bathypelagic and mesopelagic clusters consisted only of samples from those depth zones. When mesopelagic samples clustered away from other mesopelagic samples, they usually joined samples from the same sampling station (e.g., Station72 MES clusters with Station72 SRF and Station72 DCM, rather than with other mesopelagic samples). These samples were considered statistical outliers by Gregory et al. and not assigned to an ecological zone. Temperate and Tropical Epipelagic samples were dispersed among five clusters from different geographic regions. Samples also tended to cluster based on Longhurst province, except for Temperate and Tropical Mesopelagic samples.

**Figure 3.**
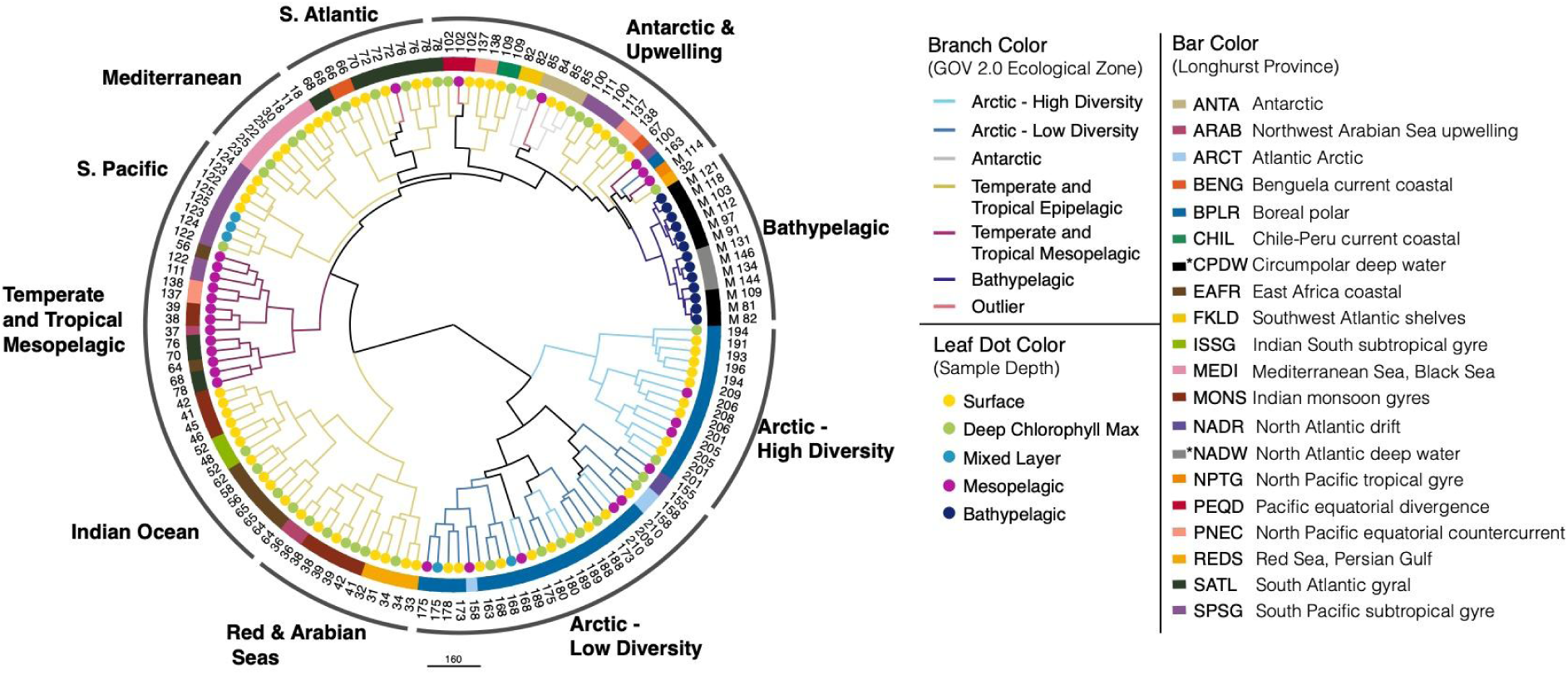
GOV 2.0 samples divide by depth and biogeography based on the composition of the RNR-encoding virus community. Samples (identified in leaf labels) were hierarchically clustered based on the abundances of all RNR-encoding viral populations in all samples using Ward’s clustering criterion as implemented in the R package hclust. The resulting dendrogram was customized in Iroki (Moore et al. 2020). Branches are colored by the ecological zones defined in the original GOV 2.0 paper. Black branches indicate that lower clades are composed of samples from different ecological regions. Leaf dots indicate sample depth and box colors indicate the Longhurst province in which the sample was taken. Asterisks in the box color legend indicate the deep water mass (CPDW and NADW) from which bathypelagic samples originated, rather than Longhurst province. Outer labels are based on the geographical region or depth from which the sample originated.

Significant differences between the RNR-encoding virus communities among sample environmental groups defined by Longhurst province, depth zone, and GOV 2.0 ecological zone were determined with perMANOVA tests, as well as individual physiochemical parameters. All environmental descriptors had a significant perMANOVA result (p < 0.05), meaning that at least one of the sample groupings was characterized by a significantly different community (Figure 4). Longhurst province explained the most variation in community structure with an R^2^ value of 0.12, followed by ecological zone (R^2^ = 0.053), and depth zone (R^2^ = 0.018).

**Figure 4.**
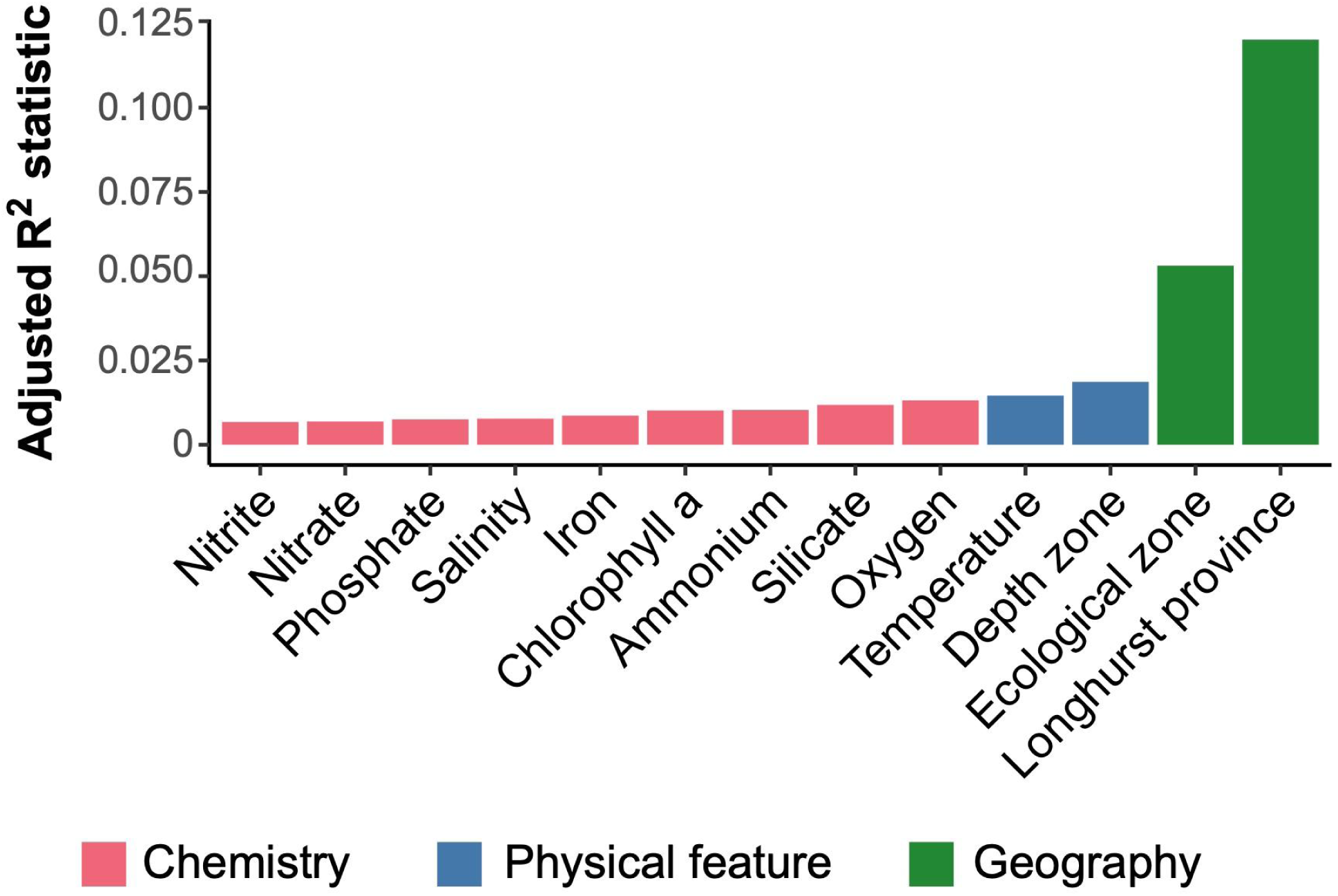
Geographical region is the best explainer of variation within the RNR-encoding community structure, with more granular environmental features explaining more of the variation. perMANOVA tests were conducted with functions from the vegan package in R using a distance-based redundancy redundancy analysis (‘dbrd’) followed by a permutational test (‘anova.cc’) with 199 permutations. ‘Rsquareadjust’ was used to adjust the default R-squared value based on the number of observations and degrees of freedom. All variables were significantly correlated to the RNR-encoding virus community (p < 0.05). Bar colors indicate variable categories.

Variation partitioning was used to examine how the measured physicochemical conditions (e.g., salinity, nitrate), geographical regions (e.g., Longhurst province), and the modeled spatial structure each influenced the composition of RNR-encoding virus communities in the ocean (Figure 5). The modeled spatial structure consisted of a spatial weight matrix that was constructed to control autocorrelation in the data. The matrix described the spatial relationships between the individual samples, incorporating statistical tests (Supplementary Figure S3) and data about ocean currents, upwelling zones, etc (*see Supplementary Methods*). Variation in the RNR-encoding community was best explained by the modeled spatial structure (43%), followed by geographical regions (33%) and lastly physicochemical parameters (27%) (Figure 5).

**Figure 5.**
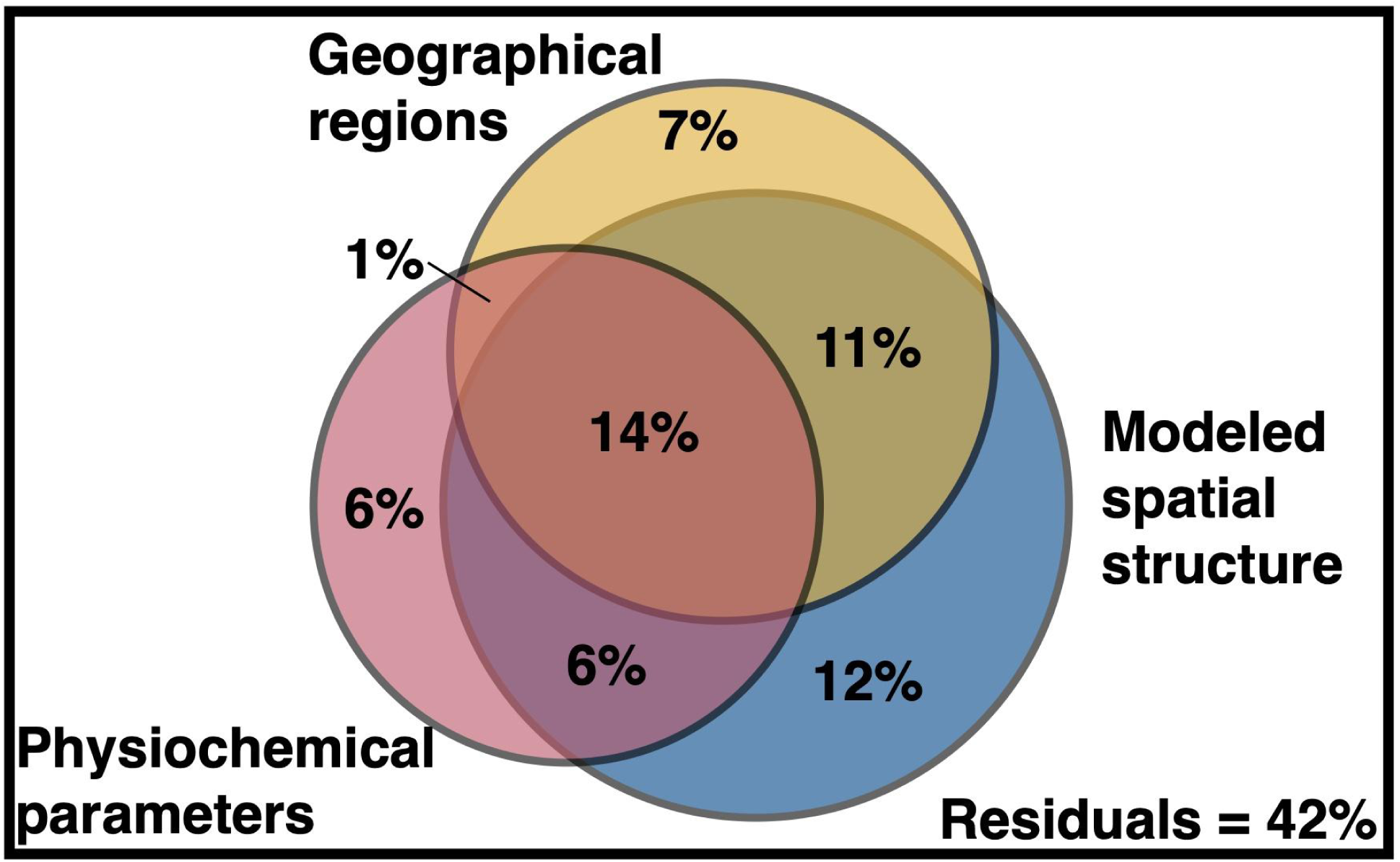
Variance partitioning reveals environmental features influencing the distribution of RNR-encoding virioplankton. Variation in the RNR-encoding virus community was partitioned among physicochemical parameters, the modeled spatial structure, and predefined geographical regions using the varpart function in R package vegan. Geographical regions include Longhurst biomes, Longhurst provinces, and the GOV 2.0 ecological zones defined in the original GOV 2.0 analysis. Individual fractions were tested for significance via redundancy analysis using the ‘rd’ and ‘anov’ functions in vegan with 999 iterations, excepting untestable fractions.

However, much of this variation could not be explained by one of these factors alone (Supplementary Figure S4); 32% of the variation was explained by some combination of the three factors. 42% of the variation was left unexplained. All partitions were significant (p ≤ 0.001).

### RNR subclass ecology

Abundances of RNR-encoding viral populations were divided based on RNR class/subclass enabling exploration of the potential role of RNRs in shaping the biology and ecology of marine viruses. Populations carrying Cyano M and Cyano SP were also analyzed as separate groups, as marine cyanophage RNRs have been previously hypothesized to have adapted to intracellular metal concentrations rather than extracellular concentrations as is hypothesized for other RNR-encoding viral populations (Harrison et al. 2019).

Viral populations containing different RNR subclasses displayed different abundance patterns with regard to iron concentrations and latitude (Figure 6). With regard to predicted iron concentrations, subclasses Ia and Id showed positive correlations, reaching a saturation point at roughly 2.5 nM. Cyano M, Cyano SP, and subclass Ic-like groups were anti-correlated with iron concentration, and Clade NrdAK and both Class II subclasses showed no relationship to predicted iron (Figure 6A).

**Figure 6.**
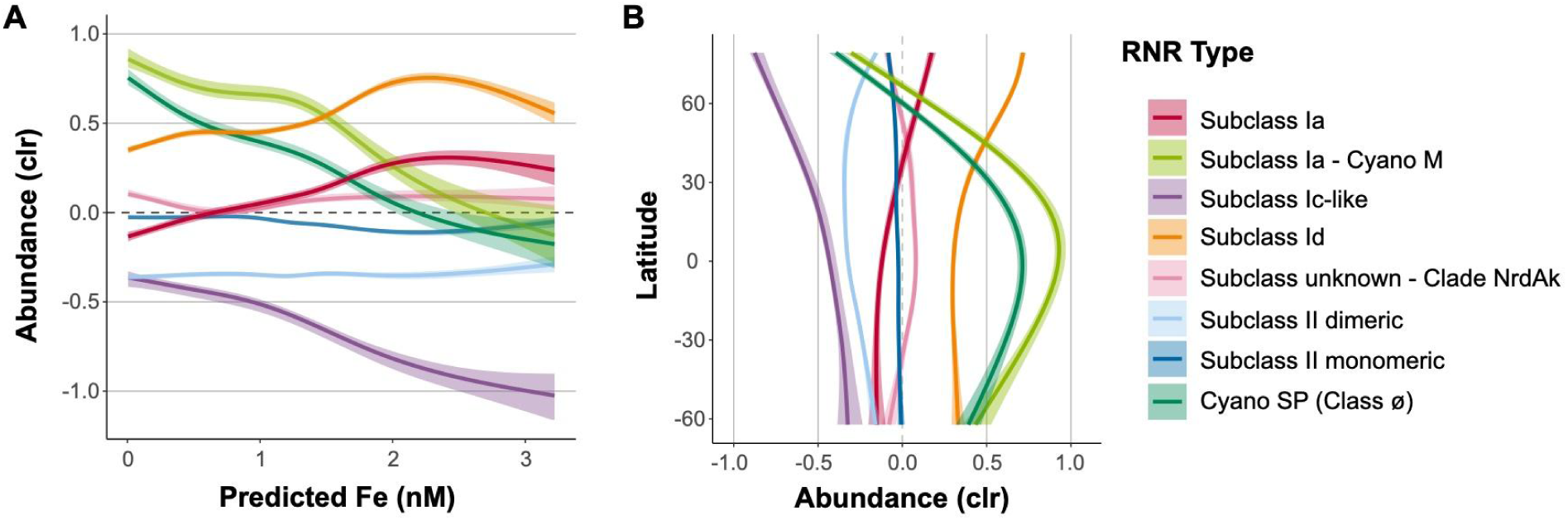
Iron concentration and geography influence distributions of RNR-encoding viral populations. Viral population abundance is divided by the type of RNR carried. A) General additive models with 95% confidence intervals depicting the relationship between CLR-transformed viral population abundance and predicted iron concentrations. Iron concentrations were predicted for epipelagic *Tara* Oceans samples using two biogeochemical models developed by a leading group of ocean trace metals researchers in conjunction with the *Tara* Oceans team (Caputi et al. 2019). Predictions shown here were generated by the PISCES model. B) General additive models with 95% confidence intervals depicting the relationship between CLR-transformed viral population abundance and latitude.

With respect to latitude, subclasses Ia and Id showed steady abundances from 60°S until around 20°N, where their abundances steadily increased, making them the most abundant groups in the Arctic (Figure 6B). Both Cyano groups peaked in the low (equatorial) latitudes and were lowest in abundance in the high latitudes. The abundances of Clade NrdAK and both Class II subclasses showed no strong relationship with latitude. The subclass Ic-like group showed its highest abundance near 60°S decreasing in abundance steadily all the way to roughly 80°N. However, samples from the northern and southern hemispheres are often not directly comparable, as samples from the northern hemisphere are predominantly from semi-enclosed and marginal seas, while samples from the southern hemisphere are largely from the open ocean (Supplementary Figure S5).

In terms of depth, subclasses Ia and Id displayed their highest abundance at the surface and their lowest abundance at the mesopelagic, with a steady decrease through the deep chlorophyll maximum (DCM) (Figure 7). The Cyano M and SP groups both showed their lowest abundance in mesopelagic waters, but Cyano SP abundance was greatest at the DCM, while Cyano M abundances were roughly equal at the DCM and the surface. The abundance of the subclass II monomeric group was seemingly unaffected by depth, while the abundance of subclass II dimeric increased greatly between the DCM and the mesopelagic. Clade NrdAk displayed similar abundance at the surface and DCM, but decreased abundance at the mesopelagic. Lastly, the subclass Ic-like group had lower concentrations at the surface and mesopelagic, but elevated abundance at the DCM. It should be noted that depth distributions of the subgroups showed the same patterns in Arctic versus non-Arctic samples, though the relative abundances changed (Supplementary Figure S6). The only exception was the Cyano SP group, which showed its greatest abundance at the surface in Arctic samples, but the DCM in non-Arctic samples.

**Figure 7.**
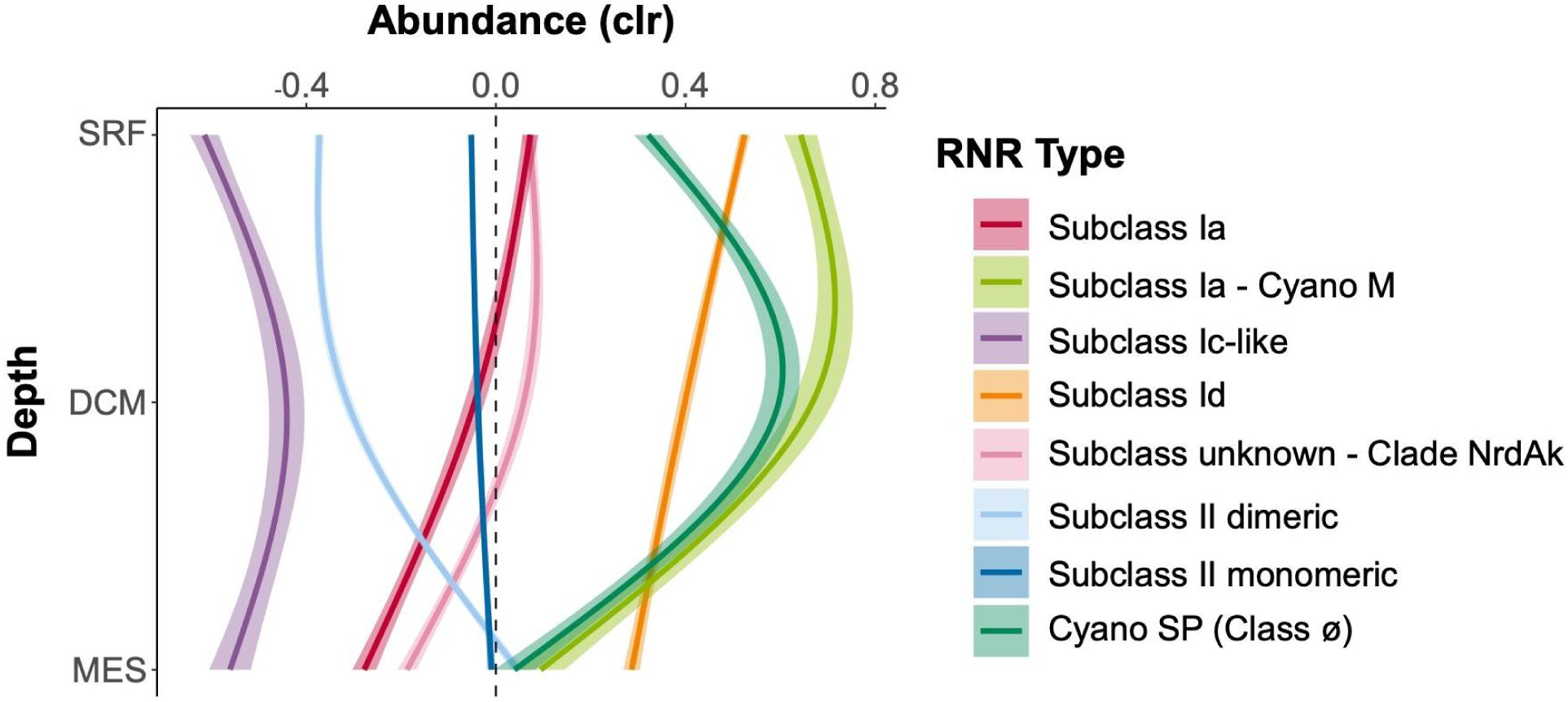
RNR-encoding viruses follow different depth distributions by RNR type. General additive models with 95% confidence intervals depicting the relationship between centered log ratio-transformed viral population abundance and depth. Viral population abundance is divided by the RNR type carried. Depths are surface (SRF), deep chlorophyll maximum (DCM), and mesopelagic (MES). Mixed layer depth and bathypelagic depths were excluded because of low sample numbers.

While the subgroups did show many different distribution patterns, there were some consistent features. In general, subclass Id or one of the Cyano groups showed the highest abundance at any depth or latitude. Clade NrdAk, contrastingly, was almost always the subgroup lowest in abundance. The Class II monomeric subclass stayed roughly at the geometric mean, showing little variation. Subclasses Ia and Id displayed highly similar patterns, but subclass Id was always more abundant. The Cyano groups also had similar patterns.

## Discussion

The RNR-encoding virus populations analyzed in this study represented nearly ten percent of all viral populations identified in the GOV 2.0 study, though the actual percentage of viruses carrying RNR genes is likely higher due to our strict validation process. A similar survey of viral RNRs has not previously been performed, so this percentage is difficult to contextualize. A past study of RNRs in marine viromes estimated that up to 93% of marine dsDNA viruses represented in RefSeq viral genomes carry RNR genes (Sakowski et al. 2014), but this was based on read mapping and abundance rather than viral populations. Another study found that RNRs were present in 17.4% of viruses surveyed (Wommack et al. 2015), but this study included few marine viruses.

### Diversity of marine viral RNRs

The viral RNRs found in the GOV 2.0 dataset represented most of the known Class I and Class II RNR subclasses. Class II RNR sequences accounted for the largest percentage of recovered sequences (∼66%), and the single largest group was the monomeric Class II RNRs (∼49%), consistent with a previous study of RNR in aquatic viromes which found that Class II monomeric RNRs made up ∼50% of RNRs (Sakowski et al. 2014). However, these observed frequencies are within previously clustered GOV 2.0 viral populations (Gregory et al. 2019), thus the actual abundance of subclass members in this study is unknown. Prior work within known viral genomes carrying RNR genes found that 24% encoded Class II RNRs, 5% Class III, and 71% Class I (Wu et al. 2023). However, as with this study, the vast majority of Class II RNRs within viral genomes were monomeric. The bias towards Class II RNRs (and in particular Class II monomeric RNRs) within the unknown marine viruses of the GOV 2.0 study is stark when comparing these findings with known viral genomes.

Class II monomeric sequences were also more phylogenetically diverse than expected. Class II monomeric sequences constitute a single small clade in the RNRdb, whereas Class I α and Class II dimeric sequences are divided among several clades (Lundin et al. 2009). The presence of several distinct Class II monomeric clades (Supplementary Figure S2) was therefore surprising. This is a strong indication that the RNR diversity in environmental datasets is not currently captured in public sequence databases which are largely composed of genome-sequenced isolates (Dwivedi et al. 2013).

RNRs belonging to the Class II dimeric subclass, Class I subclasses Ia and Id, Clade NrdAk, Cyano SP, and Cyano M were also found in many viral populations. Subclasses Ic and Ie and Class III RNRs were entirely absent in the GOV 2.0 dataset, and very few subclass Ib sequences were recovered. The sensitive homology search and the composition of the query sequences from the RNRdb make it unlikely that these sequences would have been missed. Subclasses Ib, Ic, and Ie are commonly found in host-associated, pathogenic bacteria (Blaesi et al. 2018; Johansson et al. 2010; Högbom et al. 2004), which are rare in marine ecosystems. An earlier study of viral RNRs in metagenomic datasets found an absence of subclass Ib within epipelagic marine samples, though they were abundant in a hydrothermal vent system (Dwivedi et al. 2013). Subclasses Ic and Ie were not examined in that study. Class III RNRs are oxygen sensitive (Mulliez et al. 1993), and unlikely to occur within the water column.

### RNR-encoding viruses reflect the total viral community

Our results confirmed that RNR is a useful marker for environmental phage populations. First, RNR genes appeared to be single copy in marine virus genomes. Just thirteen GOV 2.0 viral populations contained more than one RNR and no populations contained more than two RNR genes. In all cases, the two RNRs were located far apart (>40kb) on the viral population contig and were virtually identical, perhaps indicating that these contigs originated from the fusion of two highly related viral genomes, rather than a single virus with multiple RNRs. However, the incidence of a virus genome encoding more than one RNR is not unprecedented as coliphage T4 encodes both a class I and class III RNR (Logan et al. 1999; Berglund et al. 1969).

Second, the RNR-encoding virus subset reflected the viral community as a whole by recreating the same previously reported community structure despite accounting for ∼10% of all GOV 2.0 viral populations. The GOV 2.0 ecological zones characterized by different total viral communities were associated with changes in structure of the RNR-encoding viral community, as the zones mapped well onto the PCA plot (Figure 2) and dendrogram (Figure 3) and statistical testing showed that GOV 2.0 ecological zone was a significant factor in differentiating RNR-encoding virus community structure (Figure 4). Additionally, the sample clusters identified on the dendrogram (Figure 3) largely overlapped with the ecological zones defined by Gregory et al., with polar and nonpolar samples being distinct and sample depth driving separation in low and mid-latitude regions. These analyses indicate that the RNR-encoding virioplankton community reflects the total viral community. Thus, RNR is likely encoded by a broad cross-section of marine virioplankton infecting an equally broad cross-section of bacterioplankton and phytoplankton hosts.

While this may seem remarkable, this result is perhaps unsurprising when considering what is known about the diversity of RNR-encoding viruses and the GOV 2.0 dataset. Viruses that infect bacteria and archaea are overwhelmingly dsDNA viruses (Iranzo et al. 2016), with tailed phage considered to be the most diverse and abundant group (Yang et al. 2019; Brum et al. 2016; Iranzo et al. 2016; Han et al. 2024). RNRs are known to be present in many lytic dsDNA viruses, including all three families of tailed phage (previously Caudovirales) (Sakowski et al. 2014; Holmfeldt et al. 2013). The sample preparation used by all expeditions included in GOV 2.0 are biased toward dsDNA viruses smaller than 0.22 μm (Gregory et al. 2019; Roux et al. 2017), meaning that the GOV 2.0 dataset may be enriched in phages relative to large algal viruses.

Additionally, some viruses carrying RNR are considered to be ubiquitous in the oceans (Buchholz et al. 2022; Holmfeldt et al. 2013) and many infect hosts known to be highly abundant in marine ecosystems, such as members of the SAR11 and SAR116 clades (Zhang et al. 2019; Kang et al. 2013) and Cyanobacteria (Sakowski et al. 2014; Sullivan et al. 2005).

### RNR subclasses have different ecological distributions

RNR-encoding viruses showed changing biogeographical distributions based on RNR type that largely track with known distributions of RNR metal cofactors. Viral populations carrying Subclass Ia RNRs (except those from the Cyano M clade), which utilize a di-iron cofactor, and those carrying Subclass Id RNRs, which use a di-manganese cofactor, showed nearly identical ecological distributions, though Subclass Id RNR-encoding viruses were always present in higher abundance (Figure 6). These metals have similar distribution patterns (van Hulten et al. 2017; Huang et al. 2022), however, iron is more limiting than manganese (Moore et al. 2013; Browning and Moore 2023). Abundances of subclass Ia or Id RNR-encoding viruses were lower in open ocean samples (tended to be sampled in Southern Hemisphere), and more abundant in samples from semi-enclosed and marginal seas (tended to be sampled in Northern Hemisphere), exhibiting their greatest abundance in the Arctic (Figure 6B, Supplementary Figure S5). This was a similar, but not entirely identical trend to predicted iron concentrations for this dataset (Supplementary Figure S7). The coastal Arctic, Red Sea, and northern Indian Ocean have some of the highest surface iron concentrations in the global ocean (Mahowald et al. 2009; Toulza et al. 2012). In contrast, samples from the southern hemisphere were largely from upwelling zones and open ocean sites, where iron is scarcer (Mahowald et al. 2009; Toulza et al. 2012), resulting in lower abundances. With regard to depth, viruses carrying Subclass Ia and Id RNR types were found in their greatest relative abundance in surface samples, and decreased in relative abundance with depth (Figure 7). Bioavailable forms of iron and manganese follow the same pattern as the relative abundances of Subclass Ia and Id RNR-carrying viral populations, with higher concentrations at the surface than at depth. This is likely because of photochemical reactions (Grinberg et al. 2019; van Hulten et al. 2017; Sunda 2012) which convert iron and manganese to bioavailable forms in the surface ocean.

Viral populations encoding RNRs from either the Cyano M or Cyano SP clades shadowed known distributions of marine *Synechococcus* and *Prochlorococcus* (Kolberg et al. 2004; Sullivan et al. 2005; Berggren et al. 2017). Showing dramatic changes with both latitude and depth, marine *Synechococcus* and *Prochlorococcus* are most abundant within the euphotic zone at the equator and decrease in abundance with increasing latitude (Flombaum et al. 2013).

A few mesopelagic samples did show elevated abundance of Cyano M and SP RNRs (Figure 7), which could be due to the presence of low-light *Prochlorococcus* ecotypes that can be abundant below the thermocline (Thompson et al. 2018) or due to mixing events that moved cyanobacterial cells below the mixed layer. *Prochlorococcus* and other photosynthetic microorganisms have even been detected in the dark ocean, likely due to sinking (Di Cesare et al. 2020; Guo et al. 2018). This result provides evidence for the hypothesis that cyanophage Class I RNRs are unconnected to the extracellular environment, instead relying on intracellular pools of iron or manganese within their cyanobacterial hosts (Harrison et al. 2019). The distribution of their marine *Prochlorococcus* host and reliance on intracellular pools of metallic cofactors explains the anti-correlation of the relative abundances of Cyano M and Cyano SP RNR-carrying viral populations with predicted iron concentrations.

The abundance of viral populations containing Class II monomeric RNRs showed no relationship with latitude, depth, or predicted iron concentrations (Figures 6 and 7). In fact, they maintained a steady CLR-transformed abundance of roughly zero in all cases. In rare cases, B_12_ has been found to be a secondary limiting nutrient (Moore et al. 2013), but B_12_ and cobalt are generally available throughout the ocean. Viral populations containing Class II dimeric RNRs, on the other hand, did show some distribution patterns. CLR-transformed abundances of these populations increased with depth and at high latitudes corresponding with concentrations of cobalt, the metal at the center of B_12_ (Rodionov et al. 2003). Cobalt is typically depleted in non-polar surface waters due to biological uptake (Grinberg et al. 2019) and increases in concentration with depth (Tagliabue et al. 2018). In contrast, surface seawater in the Arctic contains some of the highest concentrations of cobalt in the global ocean (Tagliabue et al. 2018).

The differences between Class II monomeric and dimeric RNR-carrying varying populations was initially puzzling, as both utilize the same cofactor. However, Class II dimeric RNRs need an additional cofactor, zinc, that is not required by Class II monomeric RNRs (Loderer et al. 2017). Another distinction between mono- and dimeric Class II RNRs is their ribonucleotide substrate. Dimeric Class II RNRs reduce diphosphate ribonucleotides (NDPs), just like Class I RNRs, whereas monomeric Class II RNRs reduce triphosphate ribonucleotide substrates (NTPs) (Larsson et al. 2010; Sintchak et al. 2002). These are stark differences when considering substrate sources. Diphosphate ribonucleotides (NDPs) result from RNA degradation and thus dimeric Class II RNRs (and Class I RNRs) rely on RNA recycling and a nucleotide kinase for supplying dNTPs for DNA synthesis. In contrast, monomeric Class II RNRs access a cell’s available pool of NTPs for dNTP production. The predominance of virioplankton populations carrying Class II monomeric RNRs may reflect selective pressures for viral replication within host cells experiencing stationary phase growth, a condition generally exhibited by most bacterioplankton (Jaishankar and Srivastava 2017). Limited measurements indicate that across growth stages intercellular NTP concentrations exceed that of NDPs and are largely stable in *E. coli* cells under stationary phase growth (Buckstein et al. 2008). If this observation for *E. coli* broadly reflects nucleotide pools within heterotrophic bacterioplankton demonstrating stationary phase growth, then phages dependent on a Class II monomeric RNR would have ready access to NTP pools sufficient for genome replication.

Finally, viral populations carrying Ic-like RNRs or RNRs from Clade NrdAk showed no patterns that could be explained by the distributions of single trace metals. It is unclear if this is because they utilize mixed cofactors (e.g., Mn/Fe), another cofactor, or no metal cofactors at all (like Subclass Ie RNRs), as there are no biochemically characterized members of either group (Grinberg et al. 2019). In the case of the Ic-like RNRs, there could simply be too few observations for a pattern to appear.

### Comparisons between this study and the original GOV 2.0 analysis

The placement of individual samples into groups based on RNR composition sometimes differed between this study and the original GOV 2.0 analysis (Gregory et al. 2019). Many of these differences were observed in samples from the Benguelan, Peruvian, and Patagonian continental shelf upwelling zones. All of these locations were experiencing upwelling events during the sampling period (Ohde and Dadou 2018; Rouault et al. 2018; Valla and Piola 2015). Another study revealed that the areas of the Antarctic sampled during *Tara* Oceans are also upwelling zones (Tamsitt et al. 2017), potentially driving the clustering of Antarctic and non-polar upwelling samples (Figure 3).

Longhurst province (Longhurst 2010) was a significant factor and explained the most variation in RNR-carrying virus community structure (Figure 4). This was surprising because the original GOV 2.0 study concluded that viral assemblages were not beholden to biogeographical ocean biomes such as Longhurst province. This indicates that the RNR-encoding virus community could be more connected to shifts in environmental conditions than the total virus community, and therefore be more likely to correspond with biogeographical regions defined by nutrient characteristics, such as Longhurst provinces (Longhurst 2010). It is unclear why differences exist between the sample groups in this study and those defined by Gregory et al 2019. One possible source is the data analysis itself. Aitchison distance was used as a β-diversity metric in this study, while the previous Gregory et al. used Bray-Curtis dissimilarity. In addition to being compatible with compositional data analysis methods, Aitchison distance has also been shown to be invariant to many biases inherent in sequencing studies (McLaren et al. 2019). This makes Aitchison distance a useful choice in comparing samples from three separate survey expeditions conducted over multiple years.

This study also took steps to account for the spatial autocorrelation present in the data (Supplementary Figure S3). Spatial autocorrelation describes the phenomena in which sites that are nearer to one another also tend to have similar metadata and biological communities (Peres-Neto and Legendre 2010; Legendre 2008). In the GOV 2.0 dataset, both geographical and temporal spatial autocorrelation are present, because sites that were sampled closer together in space were also sampled closer together in time (Gregory et al. 2019; Roux et al. 2016; Brum et al. 2015; Pesant et al. 2015). Further, plankton communities were directly connected via surface ocean currents (Supplementary Figure S5), as has been previously demonstrated within this same dataset (Richter et al. 2022). This must be controlled for when testing relationships between environmental data and sequencing results. Otherwise, autocorrelation can mask significant relationships or create spurious correlations between variables (Peres-Neto and Legendre 2010; Legendre 2008).

### Conclusions

The utility of single gene surveys in environmental viral ecology has been fiercely debated (Adriaenssens and Cowan 2014; Sullivan 2015). However, examining viral populations that carry ribonucleotide reductases brings the added dimension of biochemical phenotype to the interpretation of viral population distribution patterns. Community analysis of RNR-encoding viral populations recreated many of the results from the original GOV 2.0 study (Gregory et al. 2019), but these analyses also suggest that viruses carrying RNRs may be more sensitive to local conditions, trends that may be masked in whole community analyses.

Future work is needed for determining the factors driving the abundance and activity of RNR- containing viral communities. Additional future work should include the study of viral RNRs from other habitats. In addition to the global ocean, viral RNRs have also been found in viruses from hot springs (Loderer et al. 2019), near hydrothermal vents (Dwivedi et al. 2013), in desert hypoliths (Adriaenssens et al. 2015), and in fermented foods such as kimchi (Dwivedi et al. 2013). Studying RNRs in diverse habitats and along environmental gradients would be helpful in determining how carrying RNR genes influences the biology and ecology of environmental viruses.

## Author Contributions

AH did the analysis and wrote the manuscript. RM assisted with the analysis and edited the manuscript. KW, SP, and BF contributed to study design, data interpretation, and manuscript preparation. All authors read and approved the final manuscript.

## Acknowledgments

The authors would like to thank Mark C. Keese for input on statistical analyses involving autocorrelation.

## Supplementary Materials

Supplementary Figures 1-7

Supplementary Tables 1-3 Supplementary Methods

