## Supplementary Materials for "Ribonucleotide reductases recapitulate biogeographic patterns within virioplankton according to ocean biogeochemistry"

Supplementary Figures 1-7
Supplementary Tables 1-3
Supplementary Methods

### Supplementary Figures

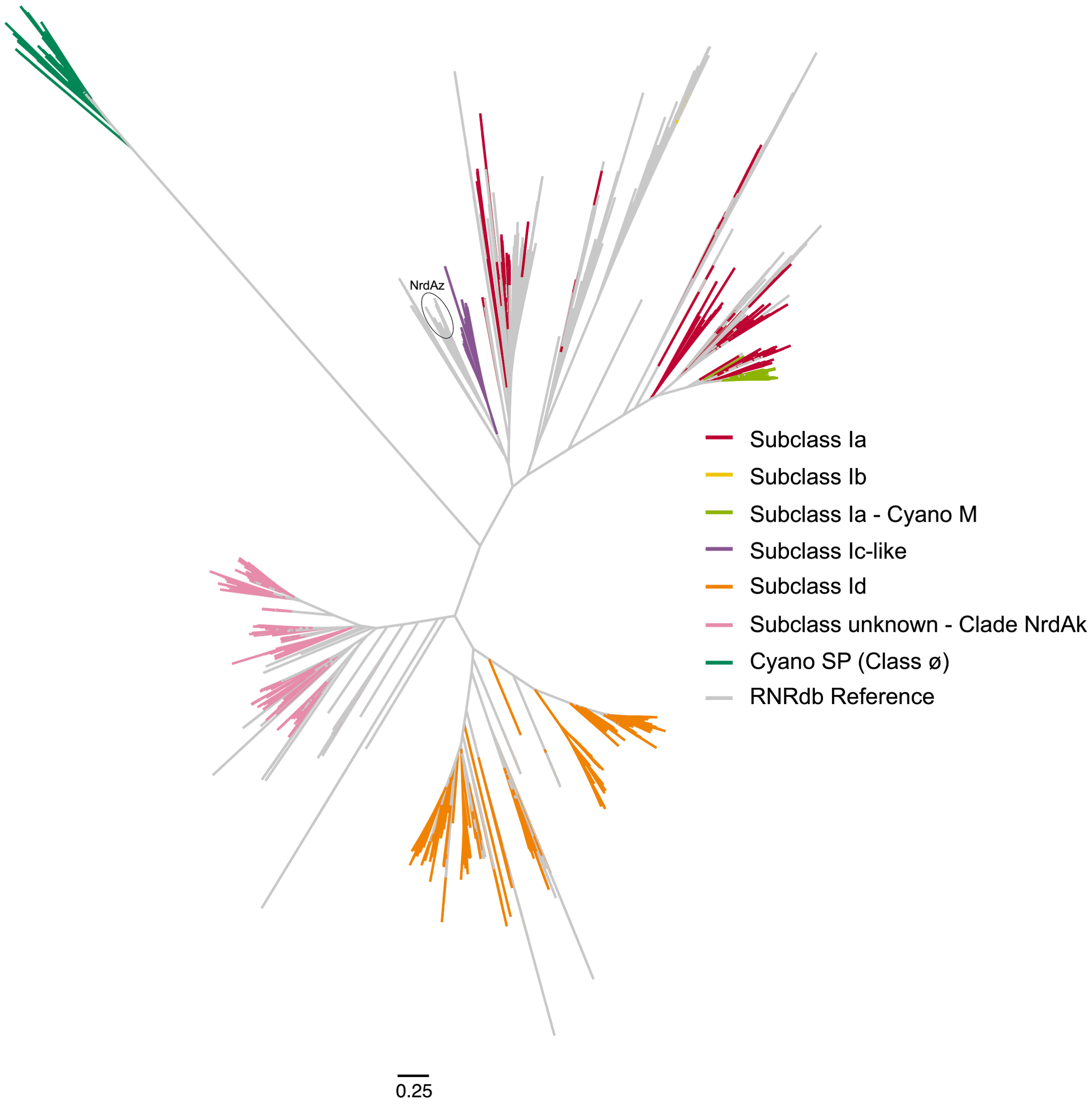


**Supplementary Figure S1.** RNR Class I and Cyano SP alpha tree used for classification. Colored branches indicate Global Ocean Virome 2.0 sequences.

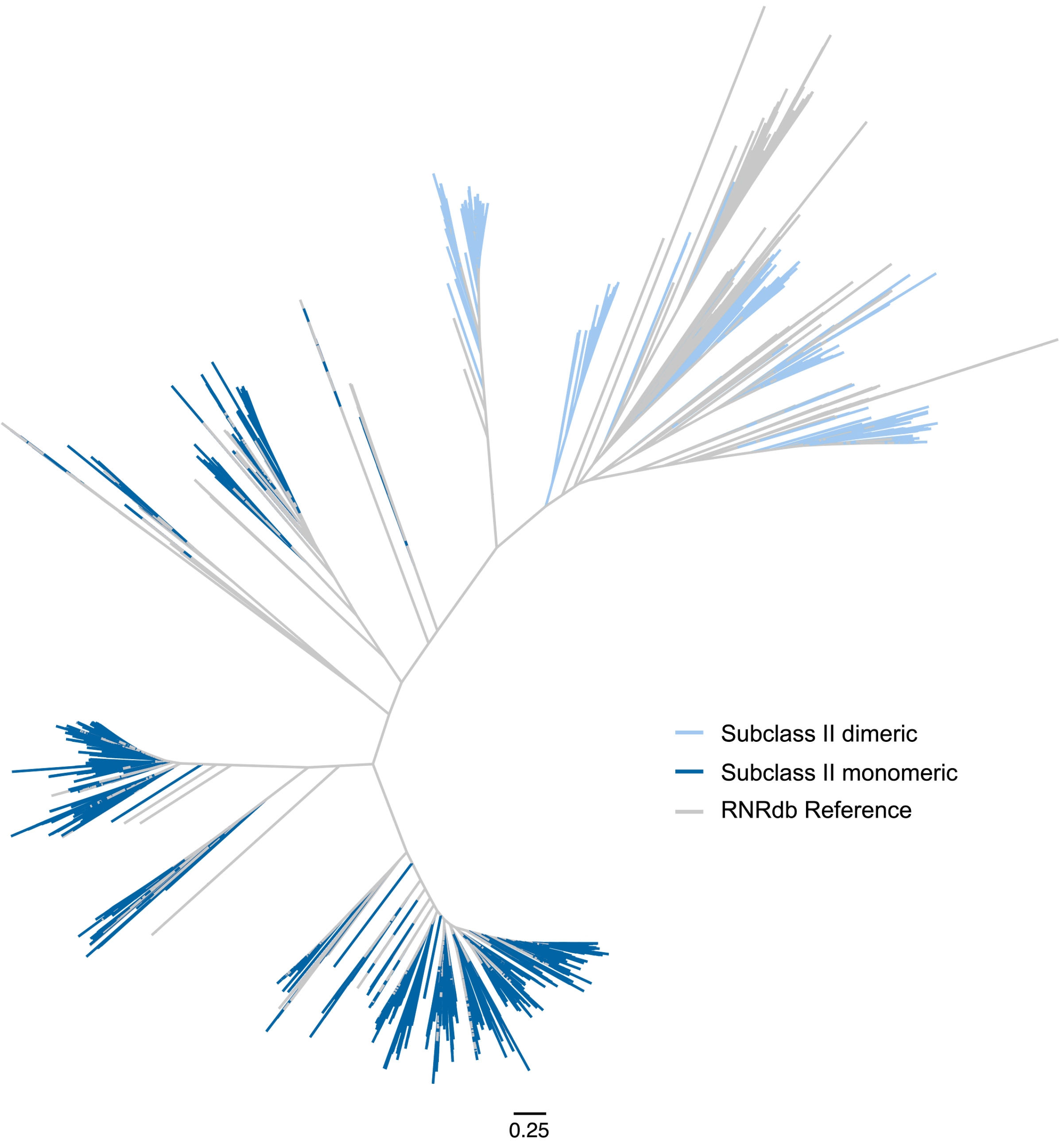


**Supplementary Figure S2.** Class II tree used for classification.

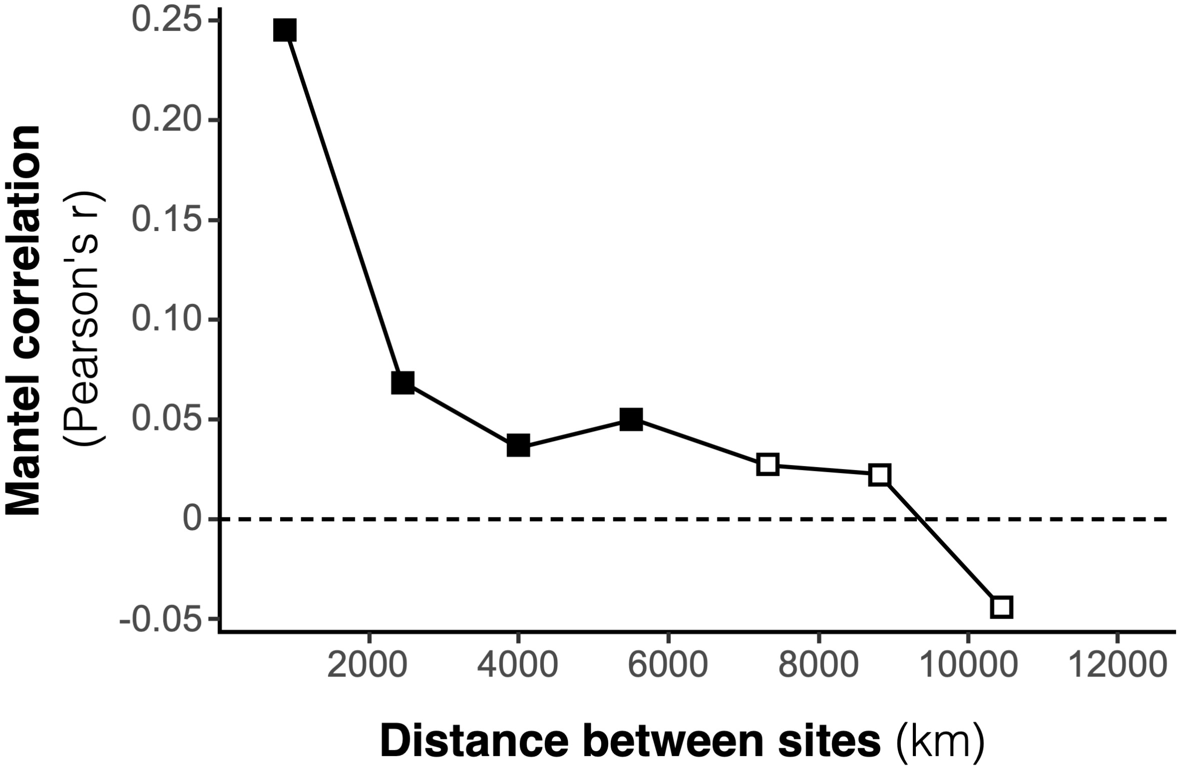


**Supplementary Figure S3.** Mantel correlogram showing Pearson's correlation between the pairwise Aitchison distance of samples and the geographical distance between samples within seven distance classes. Geographical distance between samples was calculated as the least-cost distance between samples. The Mantel correlogram was constructed using the function `mantel.correlog` from the R package `vegan`. The number of distance classes was calculated using the Sturges equation. Distance classes were tested for significant correlation using permutation Mantel tests performed by the function `mantel` from `vegan`. The Holm method was employed to correct for multiple testing.

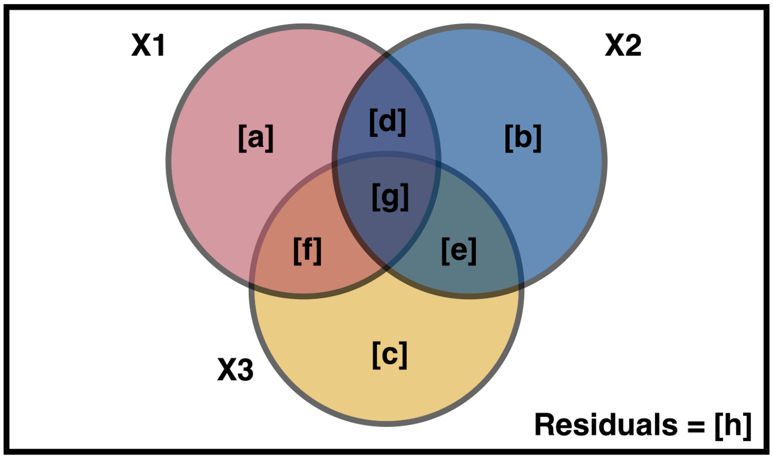


| Testable fractions: | Formula: |
|---|---|
| [a] + [b] + [c] + [d] + [e] + [f] + [g] | $Y \sim X1 + X2 + X3$ |
| [a] + [b] + [d] + [e] + [f] + [g] | $Y \sim X1 + X2 \mid X3$ |
| [a] + [c] + [d] + [e] + [f] + [g] | $Y \sim X1 + X3 \mid X2$ |
| [b] + [c] + [d] + [e] + [f] + [g] | $Y \sim X2 + X3 \mid X1$ |
| [a] + [d] + [f] + [g] | $Y \sim X1$ |
| [b] + [d] + [e] + [g] | $Y \sim X2$ |
| [c] + [e] + [f] + [g] | $Y \sim X3$ |
| [a] + [d] | $Y \sim X1 \mid X3$ |
| [a] + [f] | $Y \sim X1 \mid X2$ |
| [b] + [d] | $Y \sim X2 \mid X3$ |
| [b] + [e] | $Y \sim X2 \mid X1$ |
| [c] + [e] | $Y \sim X3 \mid X1$ |
| [c] + [f] | $Y \sim X3 \mid X2$ |
| [a] | $Y \sim X1 \mid X2 + X3$ |
| [b] | $Y \sim X2 \mid X1 + X3$ |
| [c] | $Y \sim X3 \mid X1 + X2$ |

**Supplementary Figure S4.** Explanation of variation partitioning testable fractions and formulas for significance testing, where Y is the community abundance matrix and X1, 2, and 3 are tables of explanatory variables. Plus signs (+) indicate where multiple groups of explanatory variables are being tested together. Groups of explanatory variables that are being controlled for are shown to the right of bar characters ( | ). Fractions [d], [e], [f], [g], and [h] are not testable by themselves.

**A**

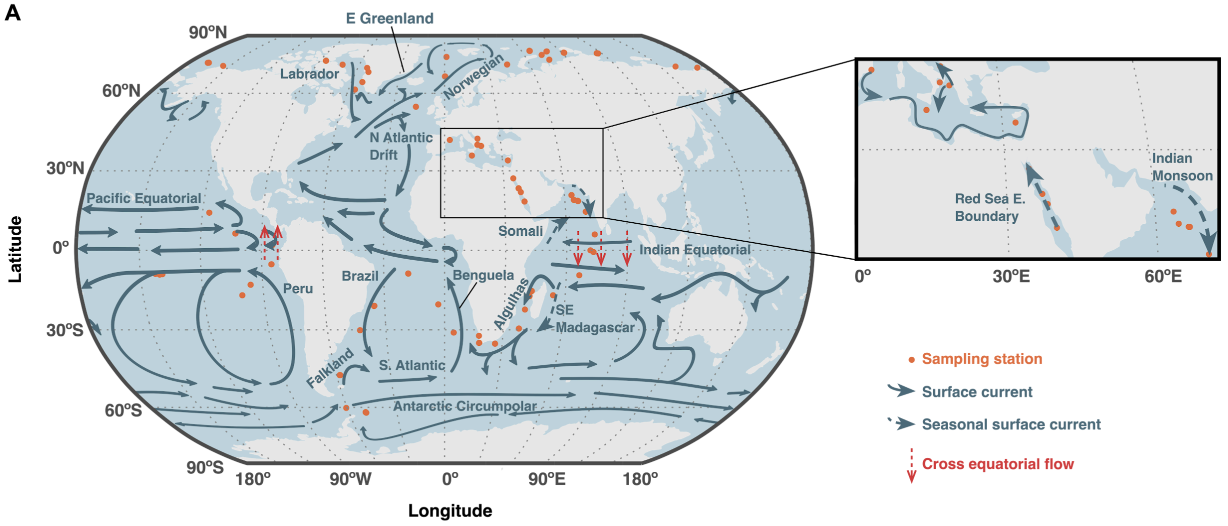


**B**

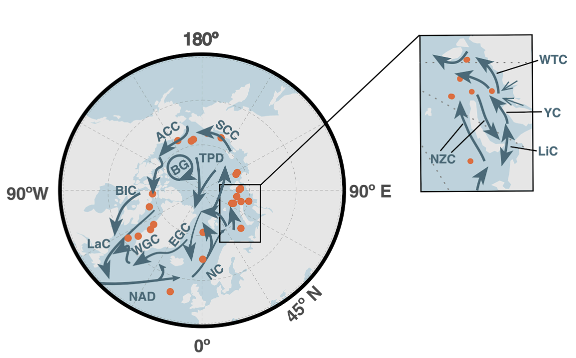


**Current Names & Abbreviations**

| | |
|---|---|
| AC | Alaskan Coastal Current |
| BIC | Baffin Island Current |
| BG | Beaufort Gyre |
| EGC | E Greenland Current |
| LaC | Labrador Current |
| LiC | Litke Current |
| NAD | North Atlantic Drift Current |
| NC | Norway Current |
| NZC | E & W Novaya Zemlya Currents |
| SCC | Siberian Coastal Current |
| TPD | Transpolar Drift Current |
| WGC | W Greenland Current |
| WTC | W Taimyr Current |
| YC | Yamal Current |

**Supplementary Figure S5.** Nearby major surface ocean currents and A) *Tara* Oceans sampling stations or B) *Tara* Oceans Polar Circle sampling stations.  Currents that alter course seasonally are shown in the orientation expected based on sampling date.

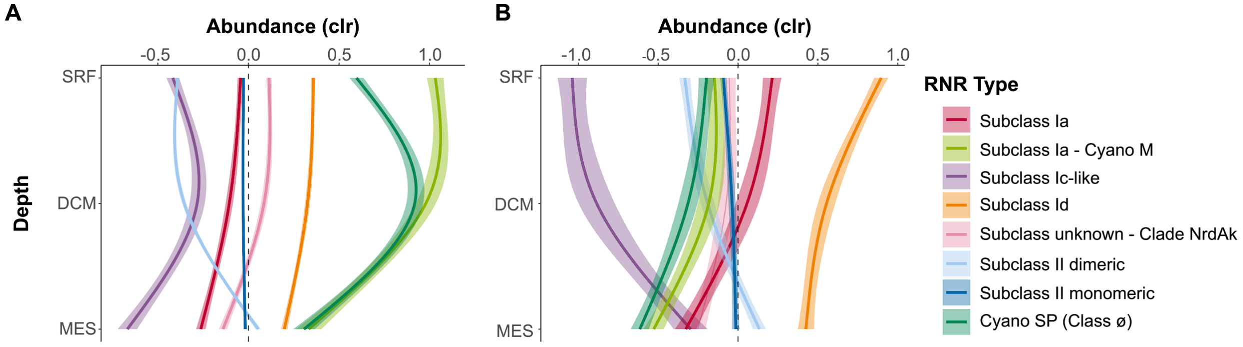


**Supplementary Figure S6.** General additive models with 95% confidence intervals depicting the relationship between centered log ratio-transformed viral population abundance and depth for A) non-Arctic and B) Arctic samples. Viral population abundance is divided by the RNR type carried. Depths are surface (SRF), deep chlorophyll maximum (DCM), and mesopelagic (MES). Mixed layer depth was excluded because of the relatively few number of samples (n = 5). The bathypelagic depth does not appear because no bathypelagic samples were taken in Arctic waters.

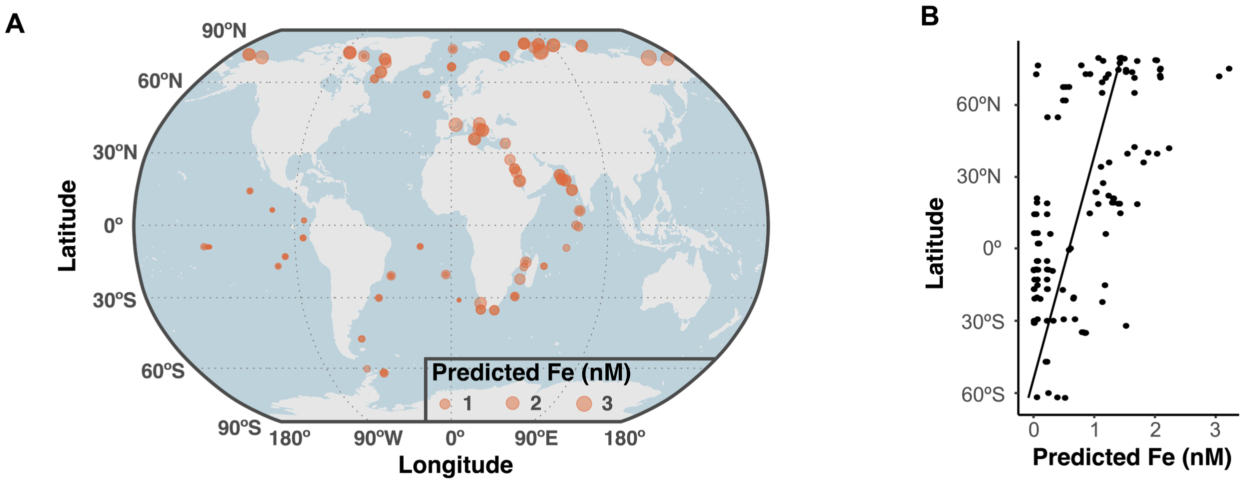


**Supplementary Figure S7.** A) Predicted iron concentrations (Caputi et al. 2019) for each *Tara* Oceans sample. B) Scatter plot of predicted iron in response to latitude with a line of best fit.

### Supplementary Tables

**Supplementary Table S1.** Conserved residues from Class I RNR alpha subunit and Class II used to validate putative sequences from GOV 2.0 contigs and their position in *E. coli*.

| Residue | Position in *E. coli* | Function | Citation |
|---|---|---|---|
| N | 437 | hydrogen bonds | Kasrayan et al. 2002 |
| C | 439 | thiyl radical - abstracts H | Mao et al. 1992a; Mao et al. 1992b |
| E | 441 | hydrogen bonds | Persson et al. 1997 |
| C | 462 | active site disulfide bridge | Mao et al. 1992a; Mao et al. 1992b |
| P | 621 | active site | Eklund et al. 2001 |

**Supplementary Table S2.** Conserved residues from Class III RNR large subunit used to validate putative sequences from GOV 2.0 contigs and their position in Bacteriophage T4.

| Residue | Position in T4 | Function | Citation |
|---|---|---|---|
| C | 79 | transient thiyl radical | Andersson et al. 2000 |
| C | 290 | transient thiyl radical - abstracts H | Olcott et al. 1998; Andersson et al. 2000 |
| C | 543 | Zn center | Andersson et al. 2000; Logan et al. 2003; Luttringer et al. 2009 |
| C | 546 | Zn center | Andersson et al. 2000; Logan et al. 2003; Luttringer et al. 2009 |
| C | 561 | Zn center | Andersson et al. 2000; Logan et al. 2003; Luttringer et al. 2009 |
| C | 564 | Zn center | Andersson et al. 2000; Logan et al. 2003; Luttringer et al. 2009 |
| G | 580 | glycyl radical | Logan et al. 1999 |

**Supplementary Table S3.** Sources for addition and manual adjustment of ocean currents in Supplementary Figure S5.

| Current | Adjustment | Citation |
|---|---|---|
| Alaskan Coastal | Addition | Stabeno et al. 2018 |
| Baffin Island | Addition | Curry et al. 2014 |
| Beaufort Gyre | Addition | Cook, n.d. |
| E &W Novaya Zemlya | Addition | Galimov et al. 2006 |
| Indian Monsoon | Addition | Yi et al. 2018; Schott et al. 2009 |
| Litke | Addition | Panteleev et al. 2007 |
| Mediterranean currents (all) | Addition | Pascual et al. 2017 |
| Red Sea E. Boundary | Addition | Taqi et al. 2019 |
| Siberian Coastal | Addition | Anderson et al. 2011 |
| Somali | Seasonal reversal | Wang et al. 2018 |
| Transpolar Drift | Addition | Constantin and Johnson 2024 |
| Western Taimyr | Addition | Panteleev et al. 2007 |
| Yamal | Addition | Galimov et al. 2006 |

### Supplementary Methods

#### GOV 2.0 Dataset

The Global Oceans Viromes (GOV) 2.0 dataset is a collection of 145 viral shotgun metagenomes taken during three expeditions (Figure 1): the Malaspina 2010 Circumnavigation (Duarte 2015), *Tara* Oceans, and *Tara* Oceans Polar Circle (TOPC) (Pesant et al. 2015). GOV 2.0 is an expansion of the original GOV dataset (Roux et al. 2016), adding the TOPC samples and additional mesopelagic samples from *Tara* Oceans.

Detailed accounts of sample collection and sequencing can be found in the publications for each of the expeditions. Briefly, Malaspina consists of 14 samples collected between April and June of 2011. The 13 bathypelagic samples were collected from 4,000 m except for two locations where ocean depth was limiting. One sample was taken from the mesopelagic (294 m) to target an oxygen minimum zone (OMZ).

The *Tara* Oceans expedition collected 90 viral samples from 45 stations between October 2009 and December 2011. The TOPC expedition collected 41 samples from 20 stations between June and December of 2013. *Tara* Oceans and TOPC samples were taken from three depths: surface water, the deep chlorophyll maximum (DCM), and the mesopelagic. All surface samples were collected at 5 m depth. The nominal depth of DCM and mesopelagic samples varies by location, as the features were identified at each site using a Rosette Vertical Sampling System (RVSS). DCM samples range from 17 to 177 m depth and mesopelagic samples range from 250 to 1,000 m depth. When no DCM was detected (stations 123, 124, and 125), samples were instead collected at the bottom of the mixed layer, which ranged from 100 to 150 m depth. Additional samples were collected when interesting environmental features were encountered, such as OMZs. All such samples were taken from the mesopelagic.

Processing and storage of samples after collection was highly similar for Malaspina, *Tara* Oceans, and TOPC samples. Following previous filtration steps, seawater (80 L, Malaspina; 20L, *Tara* and TOPC) was filtered through 0.22 μm filters. Viruses were concentrated using the iron chloride flocculation method (John et al. 2011) and viral concentrates were treated with DNase I to remove free DNA. DNA was extracted from viral concentrates using phenol/chloroform (Hurwitz et al. 2013) and sequenced on Illumina HiSeq 2000; Malaspina viromes were sequenced using 150 bp paired end protocol and *Tara* and TOPC viromes were sequenced using 101 bp paired end protocol.

Specifics of GOV 2.0 sequencing data analysis can be found in the GOV 2.0 publication (Gregory et al. 2019). Briefly, sequences underwent quality control (e.g., primer removal) and were assembled by sample of origin using metaSPAdes (Nurk et al. 2017). Assembled contigs ≥1.5 kb in length were input to VirSorter (Roux et al. 2015) and VirFinder (Ren et al. 2017) to further confirm that contigs were of viral origin. Contigs mapping to the human, cat, or dog genomes were removed from analysis. Only contigs that were verified as viral by VirSorter or VirFinder, had <40% of their length classified as cellular by the contig annotation tool (CAT) (von Meijenfeldt et al. 2019), and were ≥5 kb and linear or ≥1.5 kb and circular were used in further analysis. Remaining contigs were clustered at 95% average nucleotide identity over 80% of their length using nucmer (Kurtz et al. 2004) to create viral populations. In total, 488,130 viral populations were identified in GOV 2.0. Raw virome reads were mapped to viral populations and contigs with <5 kb coverage were removed from further analysis. Average read depth was calculated for the remaining populations and used as a proxy for abundance.

#### Spatial data Modeling

##### Testing

Malaspina samples were not included, as they were not used for statistical testing between RNR-containing viral populations and physicochemical data.

A multivariate Mantel correlogram was used to test for spatial autocorrelation in the RNR-containing viral populations (Borcard and Legendre 2012). The correlogram was calculated using the `mantel.correlog` from R package vegan v2.6-4 (Oksanen et al. 2007). Pairwise sample distances were least-cost distances computed in R using the package raster v3.4-5 (Hijmans 2023). Land was masked so that sample distance could only be calculated via ocean routes. A Moran's I test (Moran 1950) was performed for each physicochemical parameter using the function `moran.test` in R package ape v5.4-1 (Paradis and Schliep 2019) to test for significant spatial autocorrelation.

#### Spatial models

Because both the RNR-containing viral populations and physicochemical variables were significantly autocorrelated, a spatial weight matrix was constructed to control for the autocorrelation.

First, a binary connectivity matrix was manually constructed. A value of 0 or 1 was assigned to each sample pair, with 1 indicating a connection between samples. The matrix was asymmetrical to account for the directionality of ocean currents (i.e. many samples were only connected in a single direction). Samples were connected based on a few simple rules: 1) For each site pair, water was presumed to flow unidirectionally along the major currents (Supplementary Figure S5). For seasonal currents that can change direction, the time of year that sampling took place was considered in assigning directionality. 2) At some distance, sites are no longer connected. The distance limit was set as the first non-significant distance class in the Mantel correlogram (7,255,400 m; Supplementary Figure S3). 3) At the same station, water is well-mixed (in both directions) between the surface and DCM. 4) At the same station, wWater is allowed to pass from the surface and DCM to the mesopelagic, but water may not move up from the mesopelagic unless in an active upwelling zone. Sampling stations located in active upwelling zones were Station 102 - Peru upwelling zone (Boening et al. 10 2012), Station 67 - Benguela upwelling zone (Gordon et al. 1992), Station 82 - Patagonian shelf upwelling (Valla and Piola 2015), and Stations 84 and 85 - Antarctic upwelling (Baker et al. 2023; Tamsitt et al. 2017). 5) Viruses from mesopelagic samples could only be transported to other mesopelagic samples, unless in an upwelling zone. 6) Consider equatorial Ekman drift for samples separated by the equator.

The binary connectivity and least-cost distance matrices were then imported into R and the Hadamard product taken using package matrixcalc v1.0-3 (Novomestky 2022). This resulted in a matrix with least-cost distances for connected samples and zeros for unconnected samples. Based on the shape of the Mantel correlogram (Supplementary Figure S3), a spatial weight matrix was constructed by taking the inverse distance so that closer samples would be more heavily weighted than distant samples.

Because the mixed layer is a real, but leaky, barrier between surface and mesopelagic waters, a weight was used which limited the amount of water allowed to pass from surface and DCM samples to mesopelagic samples. There is no easy answer as to how much water crosses this barrier in an “average” ocean system, so we tested three additional spatial weight matrices, each with a different barrier of passage from the surface to the mesopelagic. The amount of water permitted to pass through the mixed layer was set at 50%, 10%, and 5% in each additional matrix. These three matrices, in addition to the matrix with no modeled mixed layer boundary, were compared using the function `listw.select` from R package adespatial v0.3-20 (Guénard and Legendre 2022).

Significant eigenvectors were then selected from the spatial weight matrix that explained the most variability (highest adjusted $R^2$ value) in the RNR-containing viral population (based on the centered log ratio-transformed values) for use in statistical analysis. This was done using the adespatial function `mem.select`. The spatial autocorrelation was best explained by the matrix that put no time limits on site connections and allowed 50% of viruses to cross the epi- to mesopelagic mixed layer boundary.

Taqi, Ahmed Mohammed, Abdullah Mohammed Al-Subhi, Mohammed Ali Alsaafani, and Cheriyeri Poyil Abdulla. 2019. "Estimation of Geostrophic Current in the Red Sea Based on

Sea Level Anomalies Derived from Extended Satellite Altimetry Data." *Ocean Science* 15 (3): 477–488. https://doi.org/10.5194/os-15-477-2019.

Valla, Daniel, and Alberto R. Piola. 2015. "Evidence of Upwelling Events at the Northern Patagonian Shelf Break." In *Journal of Geophysical Research: Oceans*, vol. 120, 120. no. 11. Preprint. https://doi.org/10.1002/2015jc011002.

Wang, He, Julie L. McClean, Lynne D. Talley, and Stephen Yeager. 2018. "Seasonal Cycle and Annual Reversal of the Somali Current in an Eddy‐resolving Global Ocean Model." *Journal of Geophysical Research. Oceans* 123 (9): 6562–6580. https://doi.org/10.1029/2018jc013975.

Yi, Xing, Birgit Hünicke, Nele Tim, and Eduardo Zorita. 2018. "The Relationship between Arabian Sea Upwelling and Indian Monsoon Revisited in a High Resolution Ocean Simulation." *Climate Dynamics* 50: 201–213. https://doi.org/10.1007/s00382-017-3599-8.
